# An integrative characterisation of proline *cis* and *trans* conformers in a disordered peptide

**DOI:** 10.1101/2024.05.14.594077

**Authors:** Alice J. Pettitt, Vaibhav Kumar Shukla, Angelo Miguel Figueiredo, Lydia S. Newton, Stephen McCarthy, Alethea B. Tabor, Gabriella T. Heller, Christian D. Lorenz, D. Flemming Hansen

## Abstract

Intrinsically disordered proteins (IDPs) often contain proline residues, which undergo *cis/trans* isomerisation. While molecular dynamics (MD) simulations have the potential to fully characterise the proline *cis* and *trans* sub-ensembles, they are limited by the slow timescales of isomerisation and force field inaccuracies. Nuclear magnetic resonance (NMR) spectroscopy can report on ensemble-averaged observables for both the *cis* and *trans* proline states, but a full atomistic characterisation of these sub-ensembles is challenging. Given the importance of proline *cis/trans* isomerisation for influencing the conformational sampling of disordered proteins, we employed a combination of all-atom MD simulations with enhanced sampling (metadynamics), NMR, and small-angle X-ray scattering (SAXS) to characterise the two sub-ensembles of the ORF6 C-terminal region (ORF6_CTR_) from SARS-CoV-2 corresponding to the proline-57 (P57) *cis* and *trans* states. We performed MD simulations in three distinct force fields: AMBER03ws, AMBER99SB-*disp*, and CHARMM36m, which are all optimised for disordered proteins. Each simulation was run for an accumulated time of 180-220 µs until convergence was reached, as assessed by blocking analysis. A good agreement between the *cis*-P57 populations predicted from metadynamics simulations in AMBER03ws was observed with populations obtained from experimental NMR data. Moreover, we observed good agreement between the radius of gyration predicted from the metadynamics simulations in AMBER03ws and that measured using SAXS. Our findings suggest that both the *cis*-P57 and *trans*-P57 conformations of ORF6_CTR_ are extremely dynamic and that interdisciplinary approaches combining both multi-scale computations and experiments offer avenues to explore highly dynamic states that cannot be reliably characterised by either approach in isolation.

**SIGNIFICANCE:** This study employs MD simulations (with metadynamics), NMR spectroscopy, and SAXS to elucidate the individual *cis* and *trans* proline conformations of ORF6_CTR_ from SARS-CoV-2. The good agreement on proline *cis*/*trans* populations observed in experiments (NMR) and those calculated from simulations in the AMBER03ws force field (with SAXS reweighting) showcases the efficiency of this interdisciplinary approach, which can be used to characterise highly dynamic disordered protein states, even for very slow processes. Furthermore, our study emphasises the importance of considering both computational and experimental methodologies to gain a more holistic understanding of highly dynamic proteins. The presented integrative approach sets a precedent for future studies aiming to explore complex and dynamic biological systems with slow transitions such as proline isomerisations.

## INTRODUCTION

Intrinsically disordered proteins (IDPs) and disordered regions (IDRs), which represent at least 30% of the human proteome (1), are particularly common in cancer-associated proteins, with up to 80% containing IDRs (2), and in viruses, where their coverage ranges from 3% to 55% depending on the viral species (3). Unlike folded proteins, disordered proteins are highly dynamic and they often exist as an ensemble of diverse heterogeneous conformations that lack a single three-dimensional structure. Compared to folded proteins, the primary sequences of disordered proteins have a nearly 2-fold increase of proline residues (4), which are well-known to reduce the formation of secondary structure in proteins (5). In particular, proline residues in disordered proteins have been shown to play key roles in regulating protein-protein interactions (6), posttranslational modifications (7), and liquid-liquid phase separation (8).

Most peptide bonds within proteins exist exclusively in the energetically favourable *trans* conformation. However, for proline residues, the energy barrier between the *cis* and *trans* isomers is lower due to the cyclic structure of the amino acid, approximately 85 kJ·mol^-1^ (9). Thus, isomerisation is generally a slow process, occurring at a rate of 10^-3^ - 10^-2^ s^-1^ at room temperature, depending on the adjacent residues (10,11). The *cis* proline population typically varies between 5% and 10% in disordered proteins (12), suggesting that the presence of multiple *cis* proline conformations is possible within polyproline disordered protein ensembles. These ensembles thus sample a vast conformational space of very slowly exchanging conformers, which increases the complexity (13).

Molecular dynamics (MD) simulations are often used to characterise the ensemble of disordered proteins as they can resolve individual conformations within an ensemble at atomic resolution, which is a challenge for many experimental techniques. Significant progress has been made over the last decade to optimise force fields for modelling disordered proteins (14–16), as well as advances in the integration of MD simulations and experimental data to improve their accuracy (17,18). Despite these advances, sampling the full configurational energy landscape of disordered protein ensembles in all-atom explicit solvent MD simulations is extremely computationally expensive. Proline *cis/trans* isomerisation presents an additional challenge due to the slow timescales of this process (10,11), which are generally not accessible in brute-force MD simulations alone, even on today’s most powerful computers. However, when suitable collective variables (CVs) can be identified, metadynamics, an enhanced sampling approach, offers an effective method for sampling slow motions (19,20). Indeed, metadynamics has been used to encourage exploration of the full configurational space of disordered proteins (21) and *cis/trans* proline isomerisation in simulations of dipeptides and folded systems (22,23). In the latter cases, the *ζ* angle (C^α^*_i_*_-1_, O*_i_*_-1_, C^δ^*_i_*, C^α^*_i_,* where *i* = proline) was employed as one CV for the isomerisation and pyramidalisation of the amide nitrogen (N_1_) and the *ψ* angle (N*_i_*, C^α^*_i_*, C’*_i_*, N*_i_*_+1_) was employed as an additional CV to control the amide orientation, which may affect the rate of transition between the *cis* and *trans* proline conformations. Both CVs are required to enhance proline cis/*trans* sampling as they compensate for each other.

Nuclear magnetic resonance (NMR) spectroscopy is a well-suited experimental technique to characterise ensemble-averaged properties of disordered proteins at atomic resolution under physiological conditions (pH, temperature, salt concentrations). Furthermore, NMR can uniquely characterise and quantify the populations of *cis* and *trans* proline conformations. The distinct chemical environments for the two proline isomers, coupled with their slow exchange, can result in the detection of two separate peaks for neighbouring residues or the proline itself. NMR has therefore not only been used to characterise the overall ensemble of disordered proteins (24,25), but NMR has also been used extensively to characterise the structural propensities and dynamics of *cis* proline conformations in disordered proteins (6,12,26,27).

Another experimental technique that can report on the ensembles of disordered proteins in solution is small-angle X-ray scattering (SAXS). This technique can provide coarse structural information relating to a protein’s size and shape. The capability to predict SAXS profiles from atomic coordinates, makes it possible to compare conformational ensembles from MD simulations with experimental SAXS data (28). While SAXS measurements offer powerful global information, they report ensemble-averaged states and cannot generally distinguish between the *cis* and *trans* proline configurations. Complementary approaches, such as NMR, are essential in providing detailed experimental information at the local scale.

Here, we used an integrative approach anchored in all-atom explicit solvent metadynamics simulations to characterise the C-terminal region of open reading frame 6 (ORF6_CTR_) from severe acute respiratory syndrome coronavirus 2 (SARS-CoV-2). This region of ORF6 is predicted to be disordered (Fig. S1A-B) and binds to host proteins via an essential methionine residue at position 58 (M58), leading to suppression of the innate immune response (29–31). Moreover, this 21-residue peptide contains a single proline residue at position 57 (P57), which may influence its binding to host proteins as this residue is at a preceding position to M58. We sampled the conformational space of ORF6_CTR_ using three different force fields, each optimised for disordered proteins: AMBER03ws (a03ws) (14), AMBER99SB-*disp* (a99SB-*disp*) (15), and CHARMM36m (C36m) (16). We employed metadynamics to enhance sampling (19,20), using various local and global CVs, including those on the P57 *ζ* and *ψ* angles (22,23). To reweight and validate resulting conformational ensembles, we compared ensemble-averaged properties from each force field to NMR and SAXS data. Specifically, we employed NMR chemical shifts to report on the local properties and populations of the *cis*-P57 and *trans*-P57 states, NMR diffusion experiments to compare the global properties of both states, and NMR spin-relaxation experiments to probe dynamics. Moreover, SAXS data were used to select the most accurate force field for predicting the ORF6_CTR_ global conformational ensemble. To further refine the conformational ensembles, we updated the statistical reweighting using a Bayesian/maximum entropy (BME) approach (18,32).

By integrating metadynamics simulations, SAXS, and NMR, we can characterise the highly dynamic *cis*-P57 and *trans*-P57 sub-ensembles of ORF6_CTR_. We show that metadynamics with the P57 *ζ* and *ψ* angle CVs enhances sampling of P57 isomerisation, and we observe convergence of these two CVs for the a03ws and C36m force fields. We find that a03ws most accurately predicts the *cis*-P57 and *trans*-P57 populations in the ORF6_CTR_. By employing SAXS BME reweighting (18,32) and two independent a03ws runs, we obtain *cis*-P57 populations in the a03ws force field that match those from NMR. NMR diffusion experiments indicate that the *cis*-P57 sub-ensemble is more compact than the *trans*-P57 sub-ensemble, in agreement with metadynamics simulations. Furthermore, NMR spin-relaxation experiments and metadynamics simulations suggest that both the *cis*-P57 and the *trans-*P57 conformations of ORF6_CTR_ are extremely dynamic. We anticipate that this interdisciplinary approach can be broadly applied to the many disordered proteins that undergo complex dynamics across varying timescales.

## MATERIALS AND METHODS

### ORF6_CTR_ peptide synthesis

The ORF6_CTR_ with sequence SKSLTENKYSQLDEEQPMEID was initially made by solid-phase peptide synthesis using a MultiSynTech Syro Peptide Synthesiser. Fmoc-Asp(O^t^Bu)-NovaSyn TGT resin and standard Fmoc-amino acids, coupling and deprotection conditions were used. All residues were double-coupled. The resin was cleaved using a cleavage mixture of TFA/TIPS/H_2_O (95:2.5:2.5) and the peptide was isolated by precipitation from diethyl ether, centrifugation, and lyophilisation. A portion (4 mg) of the crude peptide was purified by semi-preparative HPLC to give pure ORF6_CTR_ (1.5 mg) (see Fig. S2A-B in the supporting material).

### N-acetylated ORF6_CTR_ peptide synthesis

The unlabelled N-acetylated ORF6_CTR_ peptide (NAc-ORF6_CTR_) with sequence Ac- SKSLTENKYSQLDEEQPMEID was produced synthetically (> 96.3% purity) by GenScript (GenScript Biotech UK Limited, Oxford, UK).

### Expression and purification of the isotopically-labelled ORF6_CTR_ peptide

The uniformly isotopically labelled ORF6_CTR_ with sequence SKSLTENKYSQLDEEQPMEID was produced by recombinant protein expression with an N-terminal glutathione S-transferase (GST) tag followed by a tobacco etch virus (TEV) cleavage site in the pGEX-6P-1 expression vector (GenScript Biotech UK Limited, Oxford, UK). ORF6_CTR_ was expressed in *Escherichia coli* (*E. coli*) BL21(DE3) strain. The cells were grown at 37°C in minimal M9 media containing 1 g/L ^15^NH_4_Cl as the sole nitrogen source and 10 g/L glucose for the ^15^N-labelled ORF6_CTR_ peptide. For the ^13^C,^15^N-labelled ORF6_CTR_ peptide, 1 g/L ^15^NH_4_Cl was used as the sole nitrogen source and 3 g/L ^13^C-glucose as the sole carbon source. Cultures were grown at 37°C with vigorous shaking. Expression was induced at an OD_600_ of 0.5-0.6 by addition of 1 mM isopropyl β-D-thiogalactopyranoside (IPTG) and left shaking for 4 h at 37°C.

The cell pellet was collected by centrifugation and re-suspended in lysis buffer containing 50 mM Tris pH 8.0, 300 mM NaCl, 10 mM β-mercaptoethanol, and 5% glycerol. 0.25% IGEPAL and small amounts of DNAse, lysozyme, and protease inhibitor tablets (1 tablet per 50 mL) were added before the cells were lysed by sonication. The cell lysate was added to Glutathione Agarose 4B resin (Protino^TM^) and binding was allowed to occur by gently rocking the mixture for 1 h at 4°C. The gravity columns were then washed with the lysis buffer to remove any unbound proteins. ORF6_CTR_ was cleaved from the GST-tag via the addition of TEV protease overnight, shaking at 22°C. The next day, the flow-through containing the cleaved ORF6_CTR_ was concentrated by ultra-centrifugation through 1 kDa cut-off centricons (PALL, New York, USA) ultrafiltration membranes. For further purification, size-exclusion chromatography on a Superdex 75 column was carried out in NMR buffer containing 25 mM HEPES pH 6.9, 150 mM NaCl at 5°C. Fractions containing ORF6_CTR_ were pooled together and concentrated by 1 kDa cut-off centricons (PALL, New York, USA) ultrafiltration membranes. The yield of purified peptide was 1.4-1.6 mg per litre of culture medium.

### NMR Spectroscopy

Unless specified otherwise, NMR spectra were collected on uniformly ^13^C,^15^N-labelled or ^15^N-labelled ORF6_CTR_ peptide samples at concentrations of 300 µM and unlabelled NAc-ORF6_CTR_ at concentrations of 400 µM. All peptide samples were prepared in 25 mM HEPES buffer pH 6.9, 150 mM NaCl, containing 5% D_2_O, 1 mM sodium azide, and 1 mM EDTA. Before recording experiments, ORF6_CTR_ samples were boiled in the NMR tube (sample volume ∼600 µL) at 100°C for ∼3 minutes to remove proteases. NMR data were recorded at 15°C, unless stated otherwise. NMR data were acquired at three different static magnetic fields: 14.1 T (600 MHz) on a Bruker NEO spectrometer with a TXO cryoprobe, 18.8 T (800 MHz) on a Bruker Avance III HD spectrometer equipped with Z-gradient triple-resonance TCI cryoprobe, and 22.3 T (950 MHz) on a Bruker Avance NEO spectrometer equipped with a QCI-F cryoprobe.

Resonance assignments were obtained from a standard suite of double-resonance and triple-resonance experiments at a static magnetic field strength of 14.1 T. Backbone ^15^N relaxation rates, including *R*_1_, *R*_1ρ_, and heteronuclear {^1^H}-^15^N steady-state NOEs (hetNOEs), were measured at two static magnetic field strengths (14.1 T and 18.8 T). The backbone ^15^N exchange-free relaxation rates (*R*_dd_) were recorded using a previously described method (33) at a static magnetic field strength of 14.1 T. Diffusion Ordered SpectroscopY (DOSY) experiments were measured using a pseudo-3D ^1^H-^15^N HSQC type experiment at a static magnetic field strength of 22.3 T. Additional information regarding the NMR experiments can be found in the supporting material.

### NMR data analysis

All NMR spectra were processed using NMRPipe (34) and analysed with NMRFAM-SPARKY software (35) and FuDA (33). To analyse residual secondary structure for the *cis*-P57 and *trans*- P57 configurations, we used the secondary structure propensity score algorithm (36), with the ORF6_CTR_ ^1^H^α^, ^13^C^α^, and ^13^C^β^ chemical shifts as input. *R*_1_ and *R*_1ρ_ rate-constants were calculated by fitting the intensity profiles to mono-exponential decay functions with FuDA (33). The {^1^H}-^15^N hetNOEs were calculated as the ratios of the peak-intensities in the saturated and reference sub-spectra. The transverse relaxation rate-constants *R*_2_ were calculated from the *R*_1_ and *R*_1ρ_ relaxation rates (see Eq. S1-S2 in the supporting material). Peak intensities of the diffusion NMR data were obtained with FuDA (33) and diffusion coefficients were calculated (see Eq. S3 in the supporting material).

### SAXS measurements

The NAc-ORF6_CTR_ was dissolved in 25 mM HEPES pH 6.9, 150 mM NaCl and centrifuged at 5,600 x *g* for 10 minutes at 4°C to give a final concentration of 2 mg/mL (800 µM). The SAXS data was obtained on Instrument B21 at Diamond Light Source (Didcot, UK). Measurements were recorded at 37°C. Data sets of 26 frames with a frame exposure time of 1 s each were acquired. ScÅtter IV was used for buffer subtraction and data reduction, in which the 26 frames were averaged (37). The Guinier Peak Analysis function in ScÅtter IV was used to calculate the radius of gyration (*R*_g_). Furthermore, we rescaled the error bars for the SAXS intensities by a factor estimated through the Bayesian indirect Fourier transform (BIFT) (38) using the BioXTAS Raw software (39).

### Molecular dynamics simulations

All-atom metadynamics simulations were performed using GROMACS 2021.2 (40) patched with the open-source, community-developed PLUMED library version 2.7.1 (41). Simulations were setup using three different force fields and their corresponding water models: a03ws force field (14) with the TIP4P/2005 (42) water model, a99SB-*disp* (15) with the TIP4P-D water model (43), and C36m (16) with the CHARMM modified TIP3P water model (44). ORF6_CTR_ was N-acetylated for the simulations. The initial starting structure for NAc-ORF6_CTR_ was prepared as a linear peptide using PyMOL (45). Topology and coordinate files were then generated for the peptide using the heavy hydrogen flag for hydrogen mass repartitioning (46). The system was then solvated in a rhombic dodecahedron box with an initial volume of 774.75 nm^3^. Ions were added to neutralise the charge of the system and to maintain a concentration of 150 mM NaCl to match experiments. The solvated system was then minimised using the steepest descent algorithm with a target maximum force of 2000 kJ·mol^-1^·nm^-1^. The extended structure was then collapsed by running a high-temperature simulation at 600 K for 20 ns in the canonical (NVT) ensemble with a 2 fs time step. The collapsed peptide starting structure was then selected based on the 95^th^ percentile of the *R*_g_ to ensure a sufficiently large box size.

The collapsed peptide structure was then re-solvated and neutralised at a concentration of 150 mM NaCl in a new rhombic dodecahedron box with volumes between 290-380 nm^3^ and 9,600-12,200 explicit water molecules, depending on the force field. All subsequent simulations used a 5 fs time step. The system was minimised using the steepest descent algorithm with a target maximum force of 2000 kJ·mol^-1^·nm^-1^ and a 20 ns simulation at 600 K in the NVT ensemble was used to obtain 128 diverse starting structures. These 128 starting structures were taken at even intervals after 5 ns of equilibration time. Thermalisation was performed for 1250 ps at 310 K in the canonical (NVT) ensemble using the v-rescale thermostat (47). Then the density of the systems were equilibrated for 50 ns at 310 K in the isothermal-isobaric (NPT) ensemble using the Parrinello-Rahman barostat (48).

Production runs were executed in the NPT ensemble with a target pressure of 1 bar and temperature of 310 K using the Parrinello-Rahman barostat (48). LINCS constraints on all bonds were used (49). A Verlet list cut-off scheme was used for the non-bonded interactions. The van der Waals and Coulomb interactions were cut-off at 1.2 nm for all a03ws and a99SB-*disp* simulations and at 0.95 nm for C36m. Long-range electrostatic effects were treated with the particle-mesh Ewald method (50). Metadynamics was performed with the parallel-bias, well-tempered, and multiple-walkers protocols using a Gaussian deposition stride of 500 steps, an initial height of 1.2 kJ·mol^-1^, and a bias factor of 30 for 128 replicas (19–21). 9 CVs were selected to enhance conformational sampling (see Eq. S4-8; Table S1 in the supporting material). Metadynamics simulations were run for an accumulated time of 192 µs (a03ws run 1 and a03ws run 2), 181 µs (a99SB-*disp*), and 222 µs (C36m) until convergence was reached as assessed by blocking analysis for every CV (see Fig. S3A-C; Fig. S4; Eq. S9 in the supporting material) (28,51).

### Structural ensemble analysis

Statistical weights were calculated for each force field at the end of the simulation (see Eq. S10 in the supporting material). Analysis was conducted using the open-source package MDTraj (52). Chemical shift predictions were back-calculated from the simulations at each time step using CamShift (53). SAXS intensity curves were calculated for each structure using Pepsi-SAXS (54) and we followed a previously reported method to perform the fitting (32) We used the BME software to update the metadynamics weights to better match experimental SAXS data (18).

## RESULTS

### Sampling slowly exchanging proline *cis/trans* conformers with metadynamics simulations

To characterise the structural ensemble of ORF6_CTR_, all-atom explicit solvent MD simulations were set up with enhanced sampling provided by metadynamics (19,20). Given the sensitivity of simulations of disordered proteins to the force field used (55), we employed three diverse parameter sets (force fields) optimised for disordered proteins, capable of sampling both folded and disordered regions, or primarily extended conformations. These included the force fields: a03ws (14), a99SB-*disp* (15), and C36m (16). The ORF6_CTR_ peptide was N-acetylated (NAc-ORF6_CTR_) in the simulations to ensure a neutral charge at the N-terminus and to better match the peptide in its native context.

To obtain a good sampling of the highly heterogenous free energy surface (FES) adopted by the expected disordered peptide (Fig. S1A-B), we selected a variety of metadynamics CVs to explore both local and global properties of NAc-ORF6_CTR_. These CVs included: total α-helical content, total β-sheet content, the radius of gyration (*R*_g_), the number of salt bridges, the distance between the C^α^ of the first and last residue (end-to-end distance), the correlation between consecutive *ψ* dihedral angles (dihedral correlation), and the number of contacts between hydrophobic residues (21). To enhance proline *cis/trans* isomerisation (Fig. 1A) we also implemented 2 CVs on the P57 improper *ζ* dihedral angle and the P57 *ψ* dihedral angle (Fig. 1B) (see the supporting material for more information on the CVs) (22,23). Each CV for each force field was assessed for convergence by blocking analysis (see Fig. S3A-C; Eq. S9 in the supporting material) (28,51). We observed that the FES standard-error plateaus at a constant value for the CVs listed above in all three force fields, except for the P57 *ζ* dihedral angle CV in the a99SB-*disp* force field.

**Figure 1.**
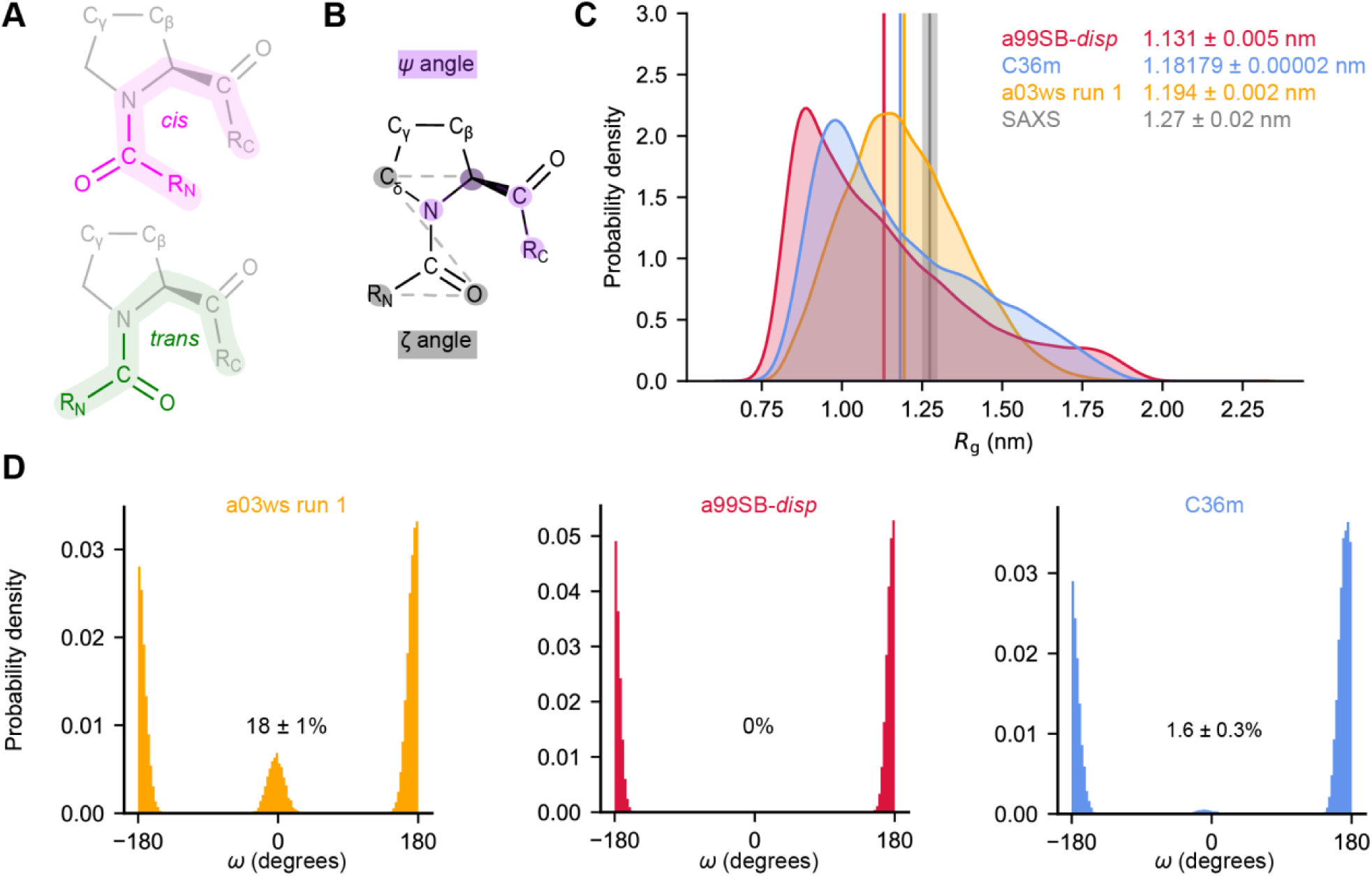
All three force fields used predict NAc-ORF6_CTR_ is disordered but only the a03ws and C36m force fields predict that NAc-ORF6_CTR_ undergoes proline *cis/trans* isomerisation. (*A*) *cis* proline (pink) and *trans* proline (green) conformations with the positions of C^γ^ and C^β^ indicated. (*B*) Definition of the *ζ* (grey) and *ψ* (purple) dihedral angles used as CVs to enhance *cis/trans* proline isomerisation in our metadynamics simulations. (*C*) *R*_g_ probability distributions were calculated using kernel density estimates to compare the a03ws run 1 (orange), a99SB-*disp* (red), and C36m (blue) ensembles. Ensemble-averaged *R*_g_ are shown for each system. The associated error represents the standard deviation between the first and second halves of the analysed trajectories. The experimental SAXS data and error (standard deviation from the Guinier analysis) are shown in grey. (*D*) Probability distribution for the *ω* dihedral angle in the a03ws run 1 (orange), a99SB-*disp* (red), and C36m (blue) ensembles, with *cis*-P57 populations displayed for each system. The error represents the standard deviation between the first and second halves of the analysed trajectories.

For the a03ws and C36m force fields, convergence was observed for the P57 *ζ* dihedral angle (Fig. S3A,C). This, combined with convergence of the *ψ* dihedral angle, suggest that P57 *cis/trans* isomerisation was converged despite the very slow timescale generally observed for proline isomerisation, typically 10 – 1000 seconds at room temperature (10). Sampling of the *cis*-P57 isomer was not observed in the simulation with the a99SB-*disp* force field (Fig. S3B). This is likely due to a99SB-*disp* hydrogen bond override, where the dihedrals were not optimised for proline and glycine (15). New and improved force fields, such as DES-Amber (56), have advanced upon this and might be better suited for proteins with both ordered and disordered regions or for investigating a disordered protein in complex with a folded protein.

To assess the sampling of secondary structures, we calculated ensemble-averaged C^α^ minimum distance contact maps for all three force fields, which showed low probabilities of α-helix and β-sheet content (Fig. S5A-C), and thus a lack of secondary structure. This result is consistent with predictions from s2D (Fig. S1A), a protein secondary structure propensity predictor trained on solution-based NMR data (57), and the AlphaFold2 pLDDT score (58), in which low confidence scores have been found to correlate with disordered regions (Fig. S1B) (59). Despite the low pLDDT scores, AlphaFold2 predicts that the ORF6_CTR_ has a helical structure, which is inconsistent with the disordered ensemble predicted in all three of our MD simulations.

### Assessing the conformational ensembles using experimental data

Histograms of the *R*_g_, calculated from the ensembles obtained with metadynamics simulations, show that NAc-ORF6_CTR_ adopts a continuum of states in all three force fields, ranging from collapsed to extended, suggesting that NAc-ORF6_CTR_ exists as an ensemble of disordered conformations (Fig. 1C) in agreement with the lack of stable secondary structure (Fig. S5A-C). It is well established that various force fields for disordered proteins can predict dramatically different features for the same disordered protein sequence, particularly in terms of ensemble-averaged properties such as *R*_g_ values (55). The three force fields used herein predict slightly different *R*_g_ values (a03ws run 1: 1.194 ± 0.002 nm, a99SB-*disp*: 1.131 ± 0.005 nm, and C36m: 1.18179 ± 0.00002 nm, Fig. 1C). To assess which of the simulations, if any, was the most in agreement with experimental measurements, we performed SAXS experiments on NAc-ORF6_CTR_ and determined an ensemble-averaged *R*_g_ of 1.27 ± 0.02 nm. Notably, a03ws and C36m performed only marginally better than the coarse-grained force field CALVADOS2, which predicts an *R*_g_ of 1.180 ± 0.007 nm (60–62). Furthermore, a recent coarse-grained IDR ensemble predictor ALBATROSS (63) predicts an *R*_g_ of 1.29 nm, which is closer in agreement with the experimental *R*_g_ than any of the all-atom models tested here. Both of these coarse-grained force field predictors specifically used the *R*_g_ parameter for optimisation (62,63).

In contrast to SAXS data, which reports on global properties, NMR chemical shifts report on local dihedral angles, ring current shifts, and electrostatics, and they are regularly used to improve and assess the accuracy of simulation ensembles because they are sensitive to secondary structure (18,21). To this end, we recorded and assigned 2D ^1^H^N^-^1^H^α^ total correlation spectroscopy (TOCSY) and ^1^H-^13^C heteronuclear single quantum coherence (HSQC) spectra of unlabelled NAc-ORF6_CTR_ to obtain ^13^C^α^, ^13^C^β^, ^1^H^α^, and ^1^H^N^ chemical shifts. Using CamShift (53), we back-calculated chemical shifts for all non-proline residues in our metadynamics ensembles for all three force fields. All experimentally-determined chemical shifts were within the CamShift error (Fig. S6A-C). To probe secondary structure in the Nac-ORF6_CTR_ we recorded a 2D ^1^H^N^-^1^H^α^TOCSY temperature titration at 8 temperature values between 5-37°C. The linear temperature coefficients of the ^1^H^N^ chemical shifts are indicative of the hydrogen bonding state of individual amides (64). For example, at higher temperatures hydrogen bonds are weakened, which causes the relative upfield shifting of the ^1^H^N^ chemical shift. All residues in the Nac-ORF6_CTR_ have temperature coefficients more negative than −4.5 ppb·K^-1^ (Fig. S7) suggesting that none of the ^1^H^N^ are involved in a long-lived hydrogen bond. This indicates that there is no substantial secondary structure in Nac-ORF6_CTR_, in agreement with the simulations.

### Solution-state NMR confirms proline *cis/trans* isomerisation

Like force field-dependent variations predicted in the *R*_g_ distributions, the *cis*-P57 population predictions also vary dramatically between different force fields (Fig. 1D). Our analysis suggests *cis*-P57 populations of 18 ± 1%, 0%, and 1.6 ± 0.3% using the a03ws run 1, a99SB-*disp*, and C36m force fields, respectively. To assess the accuracy of these *cis*-P57 populations, we recombinantly expressed and purified uniformly ^15^N-labelled and ^13^C, ^15^N-labelled ORF6_CTR_ for NMR experiments (see Materials and Methods). The production of isotopically labelled ORF6_CTR_ in *E. coli* required a GST-tag, making it extremely difficult to N-acetylate the system. While N-acetylation has been shown to decrease the *cis* proline population in small tetrapeptide systems, the effect of N-acetylation on *cis* proline sampling is reduced upon increasing the length of the peptide (12). As the N-acetylation in ORF6_CTR_ would occur at residue S41, we anticipate that potential *cis/trans* isomerisation at residue P57, separated by 16 residues from the acetylation site, is unlikely to have its chemical environment influenced by the acetyl group (see below).

Two-dimensional (2D) ^1^H-^15^N HSQC spectra show sharp resonances with a limited chemical shift dispersion in the ^1^H^N^ dimension, suggesting that these resonances arise from disordered residues (Fig. 2A). This finding agrees with spectra of the unlabelled, N-acetylated peptide (Fig. S8) and with the secondary structure predictions (Fig. S1A-B). The chemical shifts in the ORF6_CTR_ are very similar to those obtained from the NAc-ORF6_CTR_ (see Table S2 in the supporting material). The a03ws and C36m simulations discussed above suggest that the ORF6_CTR_ conformational ensemble undergoes *cis/trans* proline isomerisation about the Q56-P57 peptide bond. Indeed, in the 2D ^1^H-^15^N HSQC spectrum, two distinct signals with different signal intensities are observed for residues in proximity to P57 (Fig. 2A).

**Figure 2.**
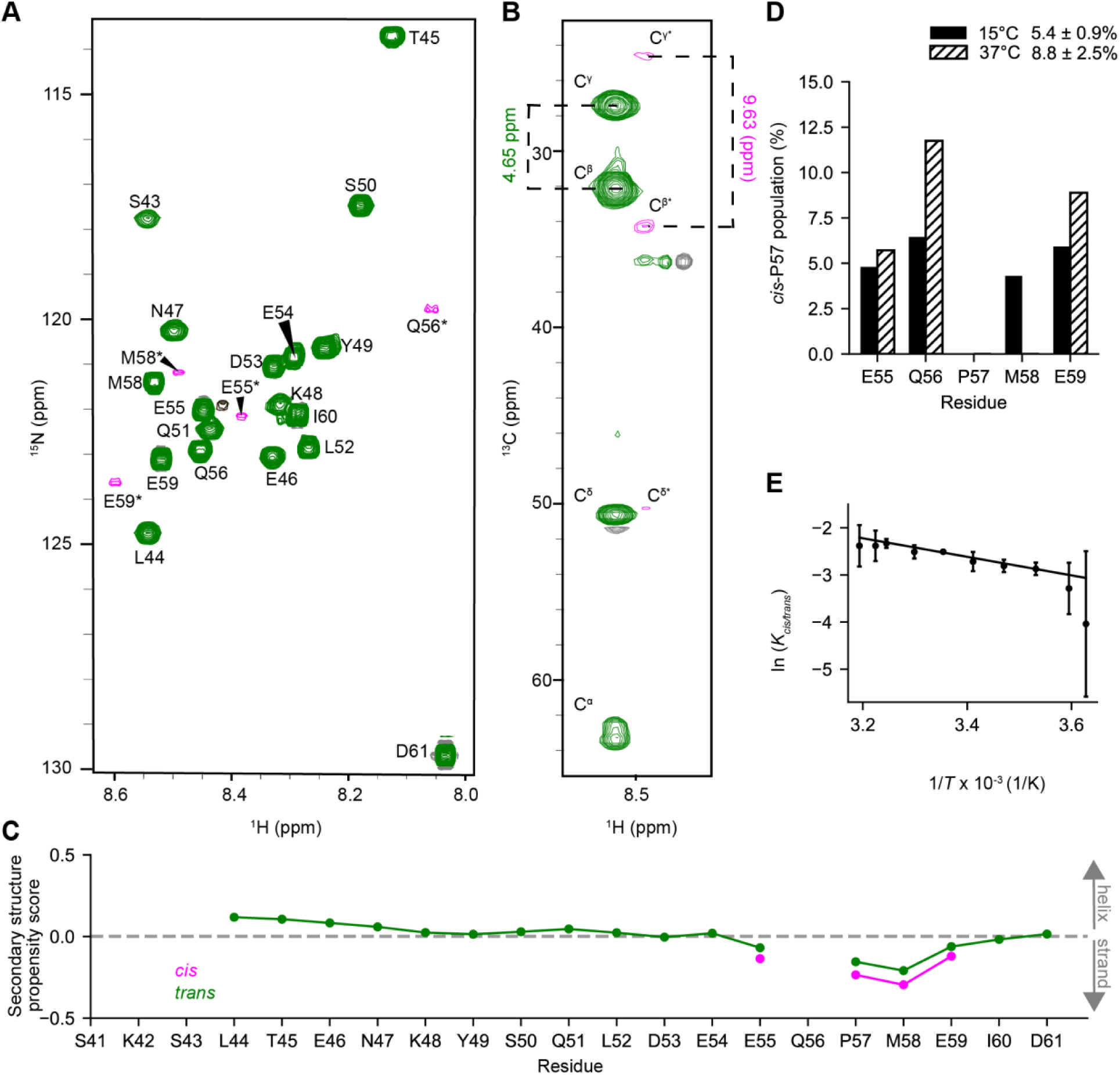
NMR verifies the disordered ORF6_CTR_ undergoes proline *cis/trans* isomerisation. (*A*) 2D ^1^H-^15^N HSQC spectrum of 300 µM ^15^N-labelled ORF6_CTR_ measured at 15°C, pH 6.9, and 14.1 T. Residues coloured in green originate from the *trans-*P57 conformational ensemble, while residues coloured in pink and labelled with an asterisk (*) report on the *cis*-P57 conformation. The black peak is unassigned. (*B*) 2D ^1^H-^13^C strip taken from the CC(CO)NH spectrum of the 300 µM ^13^C,^15^N-labelled ORF6_CTR_ at the ^15^N chemical shift of M58*. A difference of 4.65 ppm between ^13^C^β^ and ^13^C^γ^ suggests the *trans*-P57 conformation, whereas the difference of 9.63 ppm indicates the *cis*-P57 conformation. (*C*) The ORF6_CTR_ ^13^C^α^, ^13^C^β^, and ^1^H^α^ chemical shifts were used to calculate the secondary structure propensity scores for both the *cis*-P57 and *trans*-P57 sub-ensembles. A positive value indicates a helical propensity, a negative value indicates a β-strand propensity, whereas a value near zero indicates random coil. (*D*) The *cis*-P57 populations at 15°C (solid fill) and 37°C (dashed fill) are shown for each well-resolved *cis*-P57 and *trans-* P57 peak in the 2D ^1^H-^15^N HSQC spectrum. The integrated peak volume was used to calculate the mean *cis*-P57 population and standard deviation across residues E55, Q56, and E59. (*E*) The Van’t Hoff analysis of *cis/trans* P57 isomerisation determined from the integrated peak volume. The mean natural logarithm of the equilibrium constant for proline *cis*/*trans* isomerisation around the Q56-P57 bond as a function of temperature was calculated using residues E55, Q56, and E59. Error bars represent the standard deviation across the 3 residues used in the analysis. The Van’t Hoff linear fit yielded a *cis*-P57 population of 10 ± 2% at 37°C.

To verify that the additional set of minor peaks corresponds to residues in the *cis*-P57 conformational ensemble we analysed the ^13^C side-chain chemical shifts of P57 using a 3D CC(CO)NH experiment (65) (Fig. 2B). The difference in chemical shifts between ^13^C^β^ and ^13^C^γ^ is highly diagnostic of a *cis* proline (∼9.5 ppm) or a *trans* proline (∼4.5 ppm) peptide bond conformation (26,66). Similarly, peaks from the *cis*-P57 ensemble were detected for residues E55, Q56, M58, and E59. Most *cis-*P57 peaks are relatively close to their corresponding *trans-*P57 peak. For Q56, the residue preceding P57, the change in the ^15^N chemical shift between the *cis* and *trans* states is ∼3 ppm. A large change in chemical shift between *cis* and *trans* proline configurations is not uncommon for peaks of residues preceding the proline undergoing isomerisation (66). Thus, NMR provides evidence of a *cis*-P57 conformation, while also enabling the characterisation of the properties of both the proline *cis* and *trans* conformational sub-ensembles. Initially, to characterise the secondary structure content of the *cis-*P57 and *trans-*P57 sub-ensembles, we calculated the secondary structure propensity scores for the two configurations (36). These scores indicate that both conformations are disordered (Fig. 2C), although there is a slight increase in β-strand propensity near the C-terminus for the *cis-*P57 sub-ensemble.

In the slow-exchange regime, NMR peak intensities are proportional to the concentration of the species giving rise to the observed peaks (67). The population of the *cis-*P57 and *trans-*P57 conformations can thus be determined, assuming slow exchange and similar dynamics for the two conformations. The NMR experiments described above were recorded at 15°C to reduce signal loss due to backbone amide proton exchange with the solvent. The *cis*-P57 and *trans*-P57 peak intensities at 15°C were calculated for residues E55, Q56, M58, and E59 using both the integrated peak volume or peak height (Fig. 2D, Fig. S9A) (26). To enhance the *cis*-P57 population, study the system at a biologically relevant temperature, and match our metadynamics simulations, the *cis*- P57 and *trans*-P57 integrated peak volumes and peak heights were also analysed at 37°C. Residue M58 was excluded from the analysis at 37°C since the two proline isomer peaks overlap at higher temperatures. The integrated peak volume analysis yielded a *cis*-P57 population of 5.4 ± 0.9% at 15°C and 8.8 ± 2.5% at 37°C (Fig. 2D), which agrees most closely with the free energy difference between the *cis*-P57 and *trans*-P57 conformations in the a03ws run 1 metadynamics simulation.

To investigate the enthalpic and entropic factors influencing the formation of the *cis*-P57 state, we measured the equilibrium constant K = [*cis*]/[*trans*] for residues E55, Q56, and E59 at 10 temperature values between 2.5 and 40°C. From the Van’t Hoff analysis, using integrated peak volumes, we observed that the *cis*-P57 conformation is enthalpically disfavoured (Δ*Η* = 16 ± 2 kJ·mol^-1^) but entropically favoured (Δ*S* = 34 ± 7 J·K·mol^-1^) over the *trans*-P57 conformation (Fig. 2E). Similar results were obtained using peak heights, where Δ*Η* = 14 ± 1 kJ·mol^-1^ and Δ*S* = 25 ± 3 J·K·mol^-1^ (Fig. S9B). The *cis*-P57 population is consistent with results reported in the literature for other disordered protein systems (26) and is a reasonable population given that the Q56 residue, preceding P57 in ORF6_CTR_, is neither favourable nor unfavourable for proline isomerisation (66).

### Refining the metadynamics ensembles using Bayesian/maximum entropy

Given that the a03ws force field (a03ws run 1) produced an ensemble that agreed best with the experimental *cis*-P57 populations and SAXS data, we repeated the a03ws simulation (a03ws run 2) to assess reproducibility. To do this, and to avoid positive bias for the *cis*-P57 configuration, we randomly selected 128 frames only from the *trans*-P57 configuration within the initial a03ws run 1 ensemble. We then repeated the metadynamics simulation until convergence was reached at an accumulated simulation time of 192 µs (Fig. S4). The a03ws run 2 conformational ensemble was again consistent with the NMR data (Fig. S10), gave a *cis*-P57 population of 13 ± 1% (Fig. S11A) and agreed with the characterisations from a03ws run 1 (Fig. 1C; Fig. S5A; Fig. S11B-C). The robust sampling of the *cis*-P57 conformation, even when using starting structures with 100% *trans*- P57, and the consistency with both SAXS and chemical shift data from two separate metadynamics simulations, indicate that the a03ws force field produces an NAc-ORF6_CTR_ ensemble that is both robust and in agreement with the experimental data for this system.

To further refine the conformational ensemble with minimal perturbation, we used Bayesian/Maximum Entropy (BME) reweighting with the SAXS data for the two independent a03ws ensembles and the C36m ensemble (Fig. 3A; S12A) (18,32). Reweighting yielded the best χ_red_^2^ for the a03ws run 1 ensemble (Fig. 3A), but reweighting was also effective for the a03ws run 2 and C36m ensembles (Fig. S12A). As expected, reweighting resulted in a more extended ensemble-averaged *R*_g_ for the a03ws run 1 (1.292 ± 0.001 nm) and the a03ws run 2 (1.295 ± 0.005 nm) ensembles (Fig. 3B), indicating that the reweighting was successful. Reweighting also increased the ensemble-averaged *R*_g_ of the C36m ensemble (1.299 ± 0.003 nm) and shifted the C36m *R*_g_ distribution to match the a03ws distribution in both runs (Fig. S12B-D). Notably, reweighting also decreased the *cis*-P57 population in all three systems (Fig. 3C; Fig. S12E).

**Figure 3.**
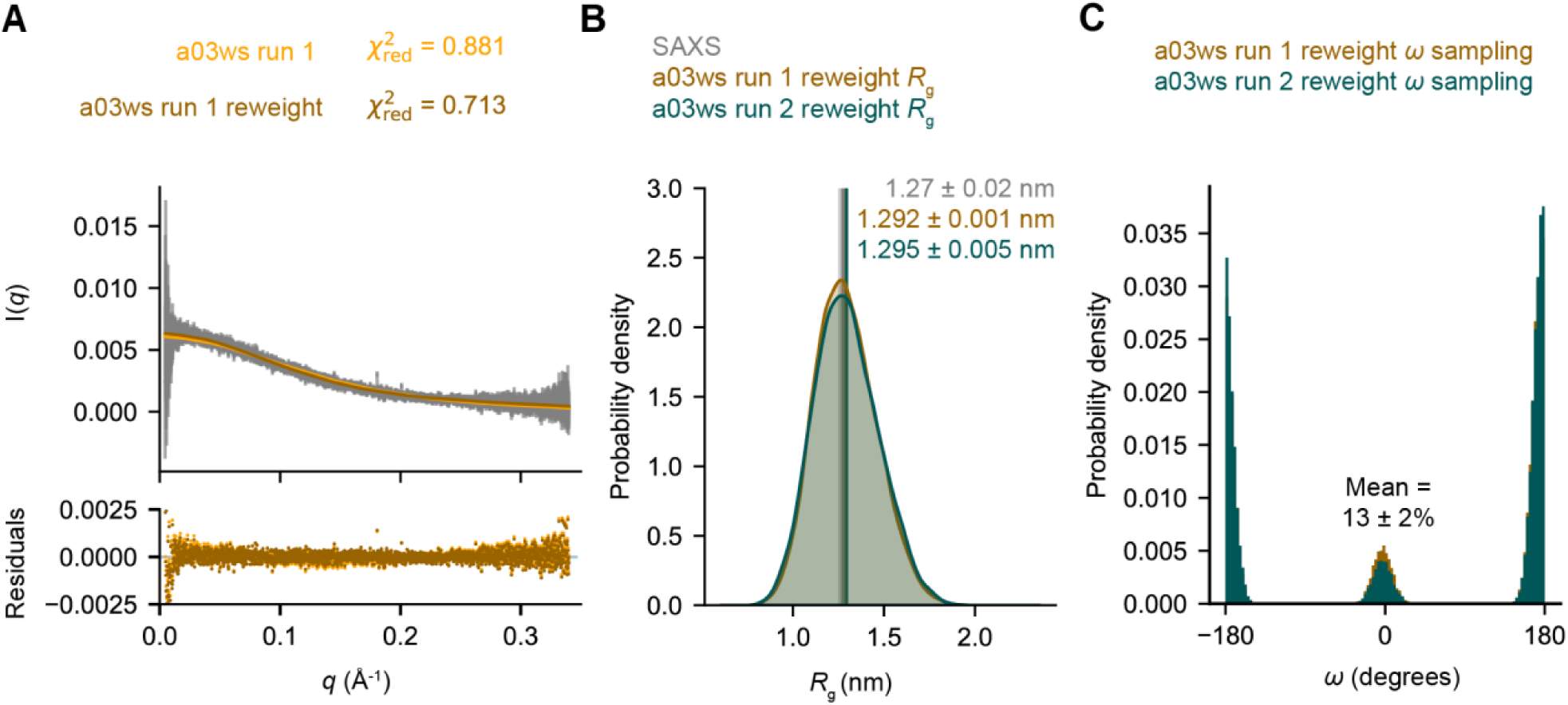
The a03ws force field most accurately predicts the disordered NAc-ORF6_CTR_ ensemble and *cis/trans* P57 populations after SAXS BME reweighting. (*A*) The calculated SAXS intensities from the a03ws run 1 metadynamics simulation (orange) and the SAXS BME reweighted simulation (brown) compared to the experimental SAXS intensities (grey), and the error associated with each intensity (grey). (*B*) *R*_g_ probability distributions were calculated using kernel density estimates to compare the SAXS BME reweighted a03ws run 1 ensemble (brown) and the SAXS BME reweighted a03ws run 2 ensemble (teal). Ensemble-averaged *R*_g_ are shown for the two runs. The associated error represents the standard deviation between the first and second halves of the analysed trajectories. The experimental SAXS data and error (standard deviation from the Guinier analysis) are shown in grey. (*C*) Probability distribution for the *ω* dihedral angle in the SAXS BME reweighted a03ws run 1 ensemble (brown) and the SAXS BME reweighted a03ws run 2 ensemble (teal). The two independent reweighted ensembles were used to calculate the mean *cis*-P57 population and standard deviation. Additional plots for the SAXS BME reweighted a03ws run 2 and C36m ensembles can be found in the supporting material.

Generally, a proline *cis* configuration within a disordered peptide induces a more compact state (13,68) and over-sampling of the *cis*-P57 conformation predicts an ensemble that is more compact than is observed from SAXS. The reweighted ensembles were compared to the chemical shift data and no significant changes were observed (Fig. S13A-C). Again, this indicates that the chemical shifts alone are not sufficient for reweighting the ensembles, as changes to the global conformation have limited influence on this parameter. Taking the two SAXS BME reweighted a03ws ensembles we calculated a mean *cis*-P57 population of 13 ± 2%, which is in good agreement with the population derived from NMR (10 ± 2%).

### ORF6_CTR_ is highly dynamic in both proline configurations

Using the reweighted metadynamics ensembles, we next sought to elucidate the structural differences between the *cis*-P57 and *trans*-P57 states. We observed that both conformations have similar *R*_g_ distributions (Fig. 4A) with the *cis-*P57 state exhibiting a slightly smaller mean conformation (1.2510 ± 0.0004 nm) as compared to the *trans*-P57 state (1.299 ± 0.003 nm). We then calculated ensemble-averaged C^α^ minimum distance contact maps for all 21 residues (Fig. 4B) and the secondary structure populations for both P57 states (Fig. 4C). Both analyses suggest an overall lack of stable secondary structure for the *cis*-P57 and the *trans*-P57 sub-ensembles. Within the N-terminal region of ORF6_CTR_, the secondary structure propensity scores calculated from NMR chemical shifts and the secondary structure populations from the metadynamics simulations indicate the sampling of very transient α-helical structures (Fig 2C; Fig. 4C; Fig. S14A-C; Fig. S15A-C). Whereas the chemical shifts around the P57 residue indicate a slight increase in β-strand. These near-miniscule changes in secondary structure were not predicted by the metadynamics simulations (Fig 2C; Fig. 4C; Fig. S14C; Fig. S15C).

**Figure 4.**
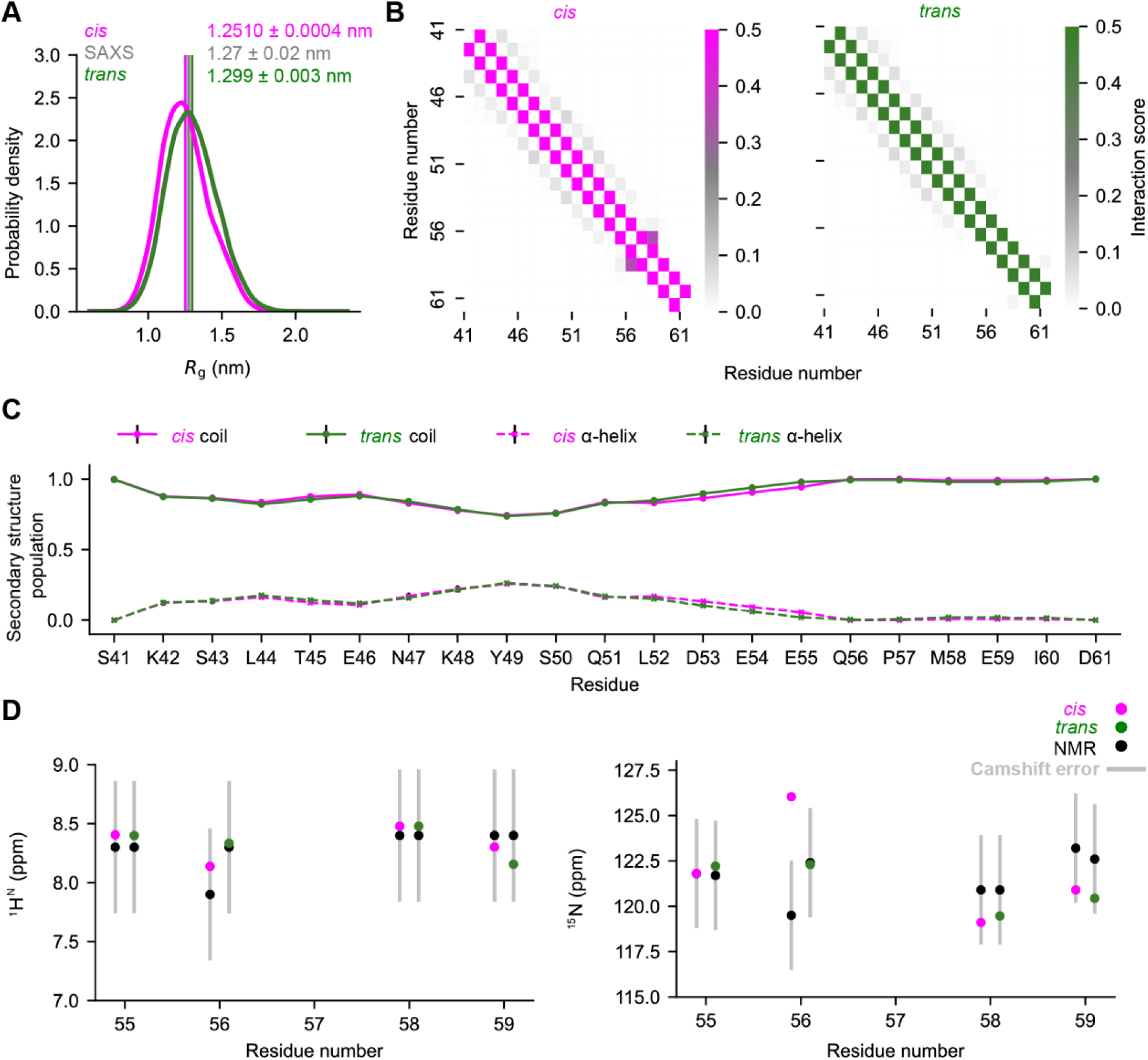
NAc-ORF6_CTR_ *cis-*P57 and *trans*-P57 sub-ensembles predicted by the a03ws run 1 SAXS BME reweighted metadynamics simulation have very similar profiles. *(A) R*_g_ probability distributions were calculated using kernel density estimates to compare the *cis*-P57 (pink) and *trans*-P57 (green) conformational sub-ensembles. Ensemble-averaged *R*_g_ are shown for each conformation. The associated error represents the standard deviation between the first and second halves of the analysed trajectory. The experimental SAXS data and error (standard deviation from the Guinier analysis) are shown in grey. (*B*) C^α^ minimum distance contact maps for the *cis*-P57 (pink) and *trans*-P57 (green) conformations. There is a low probability of α-helix and β-strand contacts in both proline configurations. (*C*) Secondary structure populations for all residues in the *cis*-P57 (pink) and the *trans*-P57 (green) conformations based on SAXS BME statistical weights. Coil populations are represented by solid lines and α-helical populations by dashed lines. β-strand represents less than 0.5% of the population for each residue so was not included. Error bars (black) represent the standard deviation between the first and second halves of the analysed trajectory. (*D*) Consistency of the ^1^H^N^ and ^15^N *cis*-P57 and *trans*-P57 ORF6_CTR_ experimental chemical shifts with the predicted chemical shifts from the SAXS BME reweighted NAc-ORF6_CTR_ a03ws run 1 ensemble. The error in CamShift (silver) is shown. The standard deviation for predicted chemical shifts between the first and second halves of the analysed trajectory were also plotted but are too small to see. See the supporting material for the a03ws run 2 and C36m analysis.

We also wondered whether the metadynamics simulations could accurately predict both the *cis*-P57 and *trans*-P57 chemical shifts using CamShift (53). Thus, we compared ^1^H^N^ and ^15^N chemical shifts for residues E55, Q56, M58, and E59 to those predicted by CamShift (Fig. 4D; Fig. S14D; Fig. S15D). The errors in CamShift are generally greater than the chemical shift differences between *cis*-P57 and *trans*-P57 states, except for the backbone nitrogen for residue Q56, which precedes P57. For this experimental observable, CamShift accurately predicts the value for the *trans*-P57 state, but inaccurately predicts that of the *cis*-P57 state (Fig. 4D; Fig. S14D; Fig. S15D). As more data for proline isomerisation of disordered proteins becomes readily available, we anticipate that force field parameters for this important state will improve, as well as the prediction of chemical shifts for *cis*-proline states.

To qualitatively compare the global features of the *cis*-P57 and *trans*-P57 ORF6_CTR_ sub-ensembles predicted by the metadynamics simulations to experiments, we recorded Diffusion Ordered Spectroscopy-HSQC (^1^H-^15^N-DOSY-HSQC) experiments at a magnetic field strength of 22.3 T. The P57 *cis*/*trans* isomerisation exchange rate is slow compared to the diffusion timescales (Δ = 200 ms), and slow compared to the longitudinal relaxation (*k*_ex_ ≪ *R*_1_), making it possible to measure distinct diffusion rates for the *cis*-P57 and *trans*-P57 sub-ensembles. We determined the average diffusion coefficients for the *cis*-P57 and *trans*-P57 conformations using data from residues Q56, M58, and E59. The diffusion measurements for the ^15^N-labelled ORF6_CTR_ at pH 6.9 and 15°C revealed a diffusion coefficient of (2.52 ± 0.09) x 10^-10^ m^2^ s^-1^ for the *cis*-P57 conformation and (2.34 ± 0.05) x 10^-10^ m^2^ s^-1^ for the *trans*-P57 conformation. The observed faster diffusion coefficient for the *cis*-P57 sub-ensemble suggests a slightly more compact conformation compared to the *trans*-P57, with statistical significance of 98% (determined via a one-tailed Welch’s t-test, df = 4, *t* = 3.179, *p* = 0.042). This finding is consistent with our SAXS BME reweighted metadynamics simulations, which predict that the ORF6_CTR_ *cis*-P57 sub-ensemble is slightly more compact than the *trans*-P57 sub-ensemble in the two a03ws and C36m simulations (Fig. 4A; Fig. S14A; Fig. S15A).

To characterise the dynamics of the ORF6_CTR_ experimentally we recorded the standard set of backbone heteronuclear ^1^H-^15^N relaxation experiments at 15°C and at two different static magnetic field strengths (14.1 T and 18.8 T) (69). ^15^N *R*_2_ rates were determined from the corresponding ^15^N *R*_1ρ_ rates, to minimise the effect of off-resonance effects and potential micro-second exchange dynamics (see Eq. S1-S2 in the supporting material) (70). The ^15^N *R*_1_ and *R*_2_ relaxation rates report on protein motions occurring at time scales faster than the effective rotational correlation time, which is usually on the nanosecond timescale for disordered proteins. The rates obtained for the ORF6_CTR_ *cis*-P57 and *trans*-P57 sub-ensembles are very similar, and both show the expected bell-shape for a random coil disordered state (Fig. 5A-B, Fig. S16A-B) (26,71).

**Figure 5.**
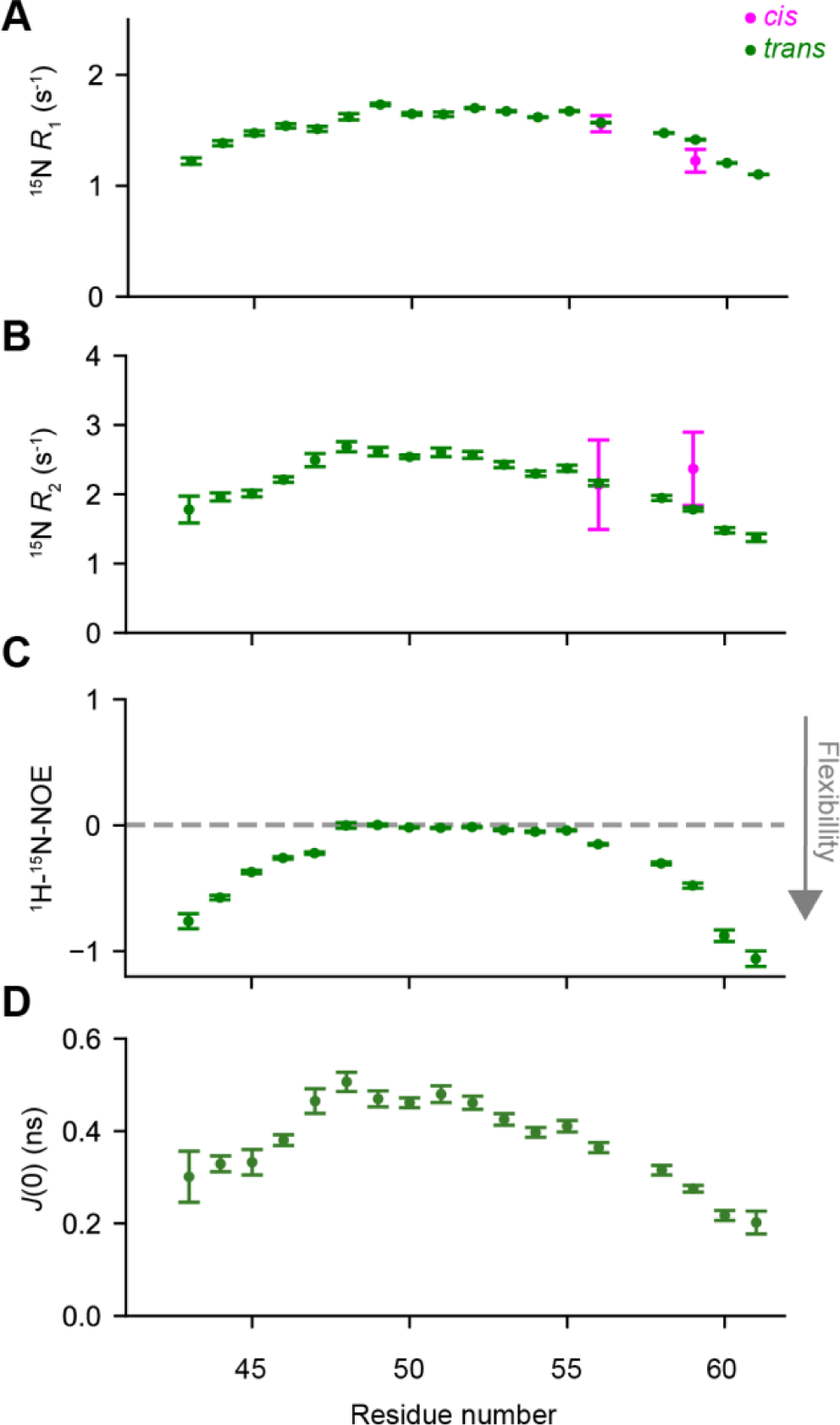
ORF6_CTR_ is very dynamic in both the *trans*-P57 and *cis*-P57 conformations. (*A*) ^15^N longitudinal relaxation rate (*R*_1_), (*B*) ^15^N transverse relaxation rate (*R*_2_), (*C*) {^1^H}-^15^N nuclear Overhauser effect (hetNOE), and (D) *J*(0) values of 300 µM ^15^N-labelled ORF6_CTR_ at 15°C and at pH 6.9. The backbone motions are shown for all of peaks corresponding to the *trans*-P57 conformation (green), and only for the well-resolved peaks with a measurable intensity corresponding to the *cis*-P57 (pink) conformation. Error bars represent the fitting of the NMR data in (*A*) and (*C*). The errors for (*B*) were calculated by error propagation. The error bars for (*D*) represent the standard deviations for each residue, which were calculated using Monte Carlo uncertainty propagation. All experiments were recorded at a static magnetic field strength of 14.1 T. {^1^H}-^15^N hetNOEs measured for the ORF6_CTR_ at a static magnetic field strengths of 14.1 T were mostly near 0 in the central region and decreased to −1 at the N- and C-termini, which is indicative of picosecond dynamics associated with a disordered protein (Fig. 5C) (69). At higher static magnetic field strengths (18.8 T) the values of the hetNOEs increase relatively, suggesting the absence of slow motions in the ORF6_CTR_ (Fig. S16C). We also evaluated the spectral density function, *J*(*ω*), at zero frequency, *J*(0), on a per-residue basis by reduced spectral density mapping (33) (Fig. 5D). This analysis indicated that both P57 sub-ensembles are highly dynamic on the nanosecond timescale and there are no substantial changes in secondary structure propensity between the two ORF6_CTR_ P57 states.

Transverse ^15^N relaxation rates (*R*_2_) also report on potential chemical exchange contribution (*R*_ex_) due to dynamics occurring on the microsecond to millisecond timescale. One way of distinguishing between the contributions to the ^15^N *R*_2_ rates arising from pico- to nanosecond dynamics and those associated with slower micro- to millisecond dynamics is through the use of the ^15^N exchange-free relaxation rate (*R*_dd_) (33). Thus, the *R*_dd_ rates were determined at a static magnetic field strength of 14.1 T (Fig. S17A), where we did not detect any exchange occurring on the microsecond to millisecond timescale (Fig. S17B-C), suggesting minimal long-range contacts and/or stabilisation of secondary structures. This is consistent with the lack of secondary structure observed for both P57 ORF6_CTR_ sub-ensembles from the secondary structure propensity scores (Fig. 2C).

## DISCUSSION

Significant progress has been made over the last decade to accurately characterise the conformational ensembles of disordered proteins with MD simulations and experimental techniques. The extensive amounts of data generated by MD simulations, alongside the capabilities of deep-learning, have now made it possible to predict some overall properties of disordered proteins from just the amino acid sequence alone (61,62). Using deep-learning techniques, global features such as the ensemble-averaged *R*_g_ can be determined within a couple of minutes. This approach achieves an accuracy that matches experimental data and surpasses the accuracy of all-atom explicit solvent MD simulations accumulated over hundreds of microseconds. The basis of these coarse-grained deep-learning approaches, have the potential to accurately characterise many global features of disordered proteins, but an atomic resolution is currently required to characterise local features such as the proline *cis* and *trans* configurations. Proline *cis*/*trans* isomerisation, which in some cases can dramatically alter the conformational ensemble of disordered proteins and thus regulate their function (6,26), is challenging to characterise with simulations given force field inaccuracies and the long simulation times required to reach convergence for this slow motion. NMR, although generally only reporting on ensemble-averaged properties, provides key insight into local features of disordered proteins, specifically proline *cis/trans* isomerisations as these conversions are slow and lead to separate NMR signals reporting on the individual proline conformations (12,26).

Using a combination of all-atom explicit solvent metadynamics simulations, and experimental NMR and SAXS, we have accurately characterised both the *cis*-P57 and *trans*-P57 conformational sub-ensembles of the 21-residue ORF6_CTR_ from SARS-CoV-2. This includes an accurate description of many of the features of both *cis-*P57 and *trans-*P57 sub-ensembles, including secondary structure content, *R*_g_, and P57 *cis*/*trans* populations. Applying the *ζ* and *ψ* CVs for P57 in ORF6_CTR_ shows that it is possible to achieve convergence for a process that generally occurs on a timescale of 10 – 1000 seconds (10,11) and in a system that is highly computationally expensive to simulate given its heterogeneous FES. While all three force fields used here produced ensembles with *R*_g_, chemical shifts predictions, and secondary structure propensities consistent with experimental data, it was only the a03ws force field that produced an ensemble that sampled a *cis-*P57 population that agreed with those calculated from NMR. The a03ws force field appears to agree best with the experimental data in the case of the very disordered peptide presented above; however, further considerations will be required if simulating other systems with distinct features. Our integrative approach will be particularly relevant for investigating the interactions of disordered protein ensembles with ligands, where proline isomerisation can be influenced by binding. The ongoing accumulation of data will further enhance the accuracy of force fields to characterise the *cis* proline state, and we anticipate that agreement between experimental ensemble-averaged proline *cis* conformations and MD resolved proline *cis* conformations will only continue to improve.

## SUPPORTING MATERIAL

Supporting material is available.

## DATA AVAILABILITY

Code that supports the findings of this study is available from GitHub at https://github.com/hansenlab-ucl/orf6-ctr_cis_trans_conformers.

## AUTHOR CONTRIBUTIONS

A.J.P. and G.T.H. performed and analysed metadynamics molecular dynamics simulations; A.J.P., V.K.S., and S.M. produced the samples; A.J.P., A.M.F., L.S.N., V.K.S., and D.F.H. performed and analysed NMR data; A.J.P. analysed SAXS data; A.B.T., G.T.H., C.D.L., and D.F.H. designed and supervised the research. All authors discussed the results and wrote the paper.

## Supporting information

orf6_str_si

## ACKNOWLEDGEMENTS

The authors acknowledge Carla Molteni, Kresten Lindorff-Larsen, Francesco Pesce, Charlie Buchanan, Carlo Camilloni, and Thomas Löhr for useful discussions. We acknowledge Greg Towers and Morten Andreas Govasli Larsen for providing the GenScript NAc-ORF6_CTR_ peptide. This work used the ARCHER2 UK National Supercomputing Service (https://www.archer2.ac.uk) under the ARCHER2 Pioneer Projects proposal (project name e692). We acknowledge PRACE for awarding us access under the special allocation COVID-19 in the PRACE Call 21, on TGCC Joliot-Curie (project name COVID1986). We acknowledge Diamond Light Source for time on beamline BL21 under proposal 32676 and Katsuaki Inoue for their assistance with the sample preparation, beamline experiments, and analysis. A.J.P. was supported by a Biotechnology and Biological Sciences Research Council UKRI funded studentship, BB/T008709/1, with the London Interdisciplinary Biosciences Consortium Doctoral Training Partnership. G.T.H. was supported by Schmidt Science Fellows, Rosaland Franklin Research Fellowship from Newnham College, Cambridge, and a BBSRC Discovery Fellowship (BB/X009955/1). L.S.N. was supported by the UCL-Birkbeck MRC DTP (MR/N013867/1). The Wellcome Trust is thanked for the award of a PhD studentship to S. M., (109073/Z/15/Z). The BBSRC (BB/R000255/1), Wellcome Trust (ref. 101569/z/13/z), and the EPSRC are acknowledged for supporting the NMR facility at University College London. Access to ultra-high field NMR spectrometers was supported by the Francis Crick Institute through provision of access to the MRC Biomedical NMR Centre. The Francis Crick Institute receives its core funding from Cancer Research UK (FC001029), the UK Medical Research Council (FC001029), and the Wellcome Trust (FC001029). For the purpose of open access, the author has applied a Creative Commons Attribution (CC BY) licence to any Author Accepted Manuscript version arising. This research is supported by the UKRI and EPSRC (EP/X036782/1).

## REFERENCES

1. Heller GT, Sormanni P, Vendruscolo M. 2015. Targeting disordered proteins with small molecules using entropy. Trends Biochem Sci. 40:491–6.

2. Iakoucheva LM, Brown CJ, Lawson JD, Obradović Z, Dunker AK. 2002. Intrinsic Disorder in Cell-signaling and Cancer-associated Proteins. J Mol Biol. 323:573–84.

3. Peng Z, Yan J, Fan X, Mizianty MJ, Xue B, Wang K, Hu G, Uversky VN, Kurgan L. 2014. Exceptionally abundant exceptions: Comprehensive characterization of intrinsic disorder in all domains of life. Cellular and Molecular Life Sciences. 72:137–51.

4. Theillet FX, Kalmar L, Tompa P, Han KH, Selenko P, Dunker AK, Daughdrill GW, Uversky VN. 2013. The alphabet of intrinsic disorder: I. Act like a Pro: On the abundance and roles of proline residues in intrinsically disordered proteins. Intrinsically Disord Proteins. 1:e24360.

5. Levitt M. 1978. Conformational preferences of amino acids in globular proteins. Biochemistry. 17:4277–85.

6. Theisen FF, Prestel A, Elkjær S, Leurs YHA, Morffy N, Strader LC, et al. 2024. Molecular switching in transcription through splicing and proline-isomerization regulates stress responses in plants. Nat Commun. 15:592.

7. Martin EW, Holehouse AS, Grace CR, Hughes A, Pappu R V., Mittag T. 2016. Sequence determinants of the conformational properties of an intrinsically disordered protein prior to and upon multisite phosphorylation. J Am Chem Soc. 138:15323.

8. Zhang X, Vigers M, McCarty J, Rauch JN, Fredrickson GH, Wilson MZ, Shea JE, Han S, Kosik KS. 2020. The proline-rich domain promotes Tau liquid-liquid phase separation in cells. Journal of Cell Biology. 219.

9. Cheng HN, Bovey FA. 1977. Cis-Trans equilibrium and kinetic studies of acetyl-L-proline and glycyl-L-proline. Biopolymers. 16:1465–72.

10. Grathwohl C, Wüthrich K. 1981. NMR studies of the rates of proline cis–trans isomerization in oligopeptides. Biopolymers. 20:2623–33.

11. Reimer U, Scherer G, Drewello M, Kruber S, Schutkowski M, Fischer G. 1998. Side-chain effects on peptidyl-prolyl cis/trans isomerisation. J Mol Biol. 279:449–60.

12. Alderson TR, Lee JH, Charlier C, Ying J, Bax A. 2018. Propensity for cis-Proline Formation in Unfolded Proteins. Chembiochem. 19:37–42.

13. Alcantara J, Stix R, Huang K, Connor A, East R, Jaramillo-Martinez V, Stollar EJ, Ball KA. 2021. An Unbound Proline-Rich Signaling Peptide Frequently Samples Cis Conformations in Gaussian Accelerated Molecular Dynamics Simulations. Front Mol Biosci. 8:734169.

14. Best RB, Zheng W, Mittal J. 2014. Balanced Protein–Water Interactions Improve Properties of Disordered Proteins and Non-Specific Protein Association. J Chem Theory Comput. 10:5113.

15. Robustelli P, Piana S, Shaw DE. 2018. Developing a molecular dynamics force field for both folded and disordered protein states. Proc Natl Acad Sci U S A. 115:E4758–66.

16. Huang J, Rauscher S, Nawrocki G, Ran T, Feig M, De Groot BL, Grubmüller H, MacKerell AD. 2016. CHARMM36m: an improved force field for folded and intrinsically disordered proteins. Nat Methods. 14:71–3.

17. Bonomi M, Heller GT, Camilloni C, Vendruscolo M. 2017. Principles of protein structural ensemble determination. Curr Opin Struct Biol. 42:106–16.

18. Bottaro S, Bengtsen T, Lindorff-Larsen K. 2020. Integrating Molecular Simulation and Experimental Data: A Bayesian/Maximum Entropy Reweighting Approach. Methods in molecular biology. 2112:219–40.

19. Barducci A, Bonomi M, Parrinello M. 2011. Metadynamics. Wiley Interdiscip Rev Comput Mol Sci. 1:826–43.

20. Pfaendtner J, Bonomi M. 2015. Efficient Sampling of High-Dimensional Free-Energy Landscapes with Parallel Bias Metadynamics. J Chem Theory Comput. 11:5062–7.

21. Heller GT, Aprile FA, Michaels TCT, Limbocker R, Perni M, Ruggeri FS, et al. 2020. Small-molecule sequestration of amyloid-β as a drug discovery strategy for Alzheimer’s disease. Sci Adv. 6.

22. Melis C, Bussi G, Lummis SCR, Molteni C. 2009. Trans-cis switching mechanisms in proline analogues and their relevance for the gating of the 5-HT3 receptor. J Phys Chem B. 113:12148–53.

23. Maschio MC, Fregoni J, Molteni C, Corni S. 2021. Proline isomerization effects in the amyloidogenic protein β 2 -microglobulin. Physical Chemistry Chemical Physics. 23:356– 67.

24. Jensen MR, Zweckstetter M, Huang JR, Blackledge M. 2014. Exploring free-energy landscapes of intrinsically disordered proteins at atomic resolution using NMR spectroscopy. Chem Rev. 114:6632–60.

25. Felli IC, Pierattelli R. 2014. Novel methods based on 13C detection to study intrinsically disordered proteins. Journal of Magnetic Resonance. 241:115–25.

26. Alderson TR, Benesch JLP, Baldwin AJ. 2017. Proline isomerization in the C-terminal region of HSP27. Cell Stress Chaperones. 22:639–51.

27. Mateos B, Conrad-Billroth C, Schiavina M, Beier A, Kontaxis G, Konrat R, Felli IC, Pierattelli R. 2020. The Ambivalent Role of Proline Residues in an Intrinsically Disordered Protein: From Disorder Promoters to Compaction Facilitators. J Mol Biol. 432:3093–111.

28. Pesce F, Newcombe EA, Seiffert P, Tranchant EE, Olsen JG, Grace CR, Kragelund BB, Lindorff-Larsen K. 2023. Assessment of models for calculating the hydrodynamic radius of intrinsically disordered proteins. Biophys J. 122:310–21.

29. Li T, Wen Y, Guo H, Yang T, Yang H, Ji X. 2022. Molecular Mechanism of SARS-CoVs Orf6 Targeting the Rae1–Nup98 Complex to Compete With mRNA Nuclear Export. Front Mol Biosci. 8:1345.

30. Gao X, Tian H, Zhu K, Li Q, Hao W, Wang L, Qin B, Deng H, Cui S. 2022. Structural basis for Sarbecovirus ORF6 mediated blockage of nucleocytoplasmic transport. Nat Commun. 13:4782.

31. Miorin L, Kehrer T, Sanchez-Aparicio MT, Zhang K, Cohen P, Patel RS, et al. 2020. SARS-CoV-2 Orf6 hijacks Nup98 to block STAT nuclear import and antagonize interferon signaling. Proc Natl Acad Sci U S A. 117:28344–54.

32. Ahmed MC, Skaanning LK, Jussupow A, Newcombe EA, Kragelund BB, Camilloni C, Langkilde AE, Lindorff-Larsen K. 2021. Refinement of α-Synuclein Ensembles Against SAXS Data: Comparison of Force Fields and Methods. Front Mol Biosci. 8:654333.

33. Hansen DF, Yang D, Feng H, Zhou Z, Wiesner S, Bai Y, Kay LE. 2007. An exchange-free measure of 15N transverse relaxation: an NMR spectroscopy application to the study of a folding intermediate with pervasive chemical exchange. J Am Chem Soc. 129:11468–79.

34. Delaglio F, Grzesiek S, Vuister GW, Zhu G, Pfeifer J, Bax A. 1995. NMRPipe: a multidimensional spectral processing system based on UNIX pipes. J Biomol NMR. 6:277–93.

35. Lee W, Tonelli M, Markley JL. 2015. NMRFAM-SPARKY: enhanced software for biomolecular NMR spectroscopy. Bioinformatics. 31:1325.

36. Marsh JA, Singh VK, Jia Z, Forman-Kay JD. 2006. Sensitivity of secondary structure propensities to sequence differences between α- and γ-synuclein: Implications for fibrillation. Protein Science. 15:2795.

37. [cited 2024 Jan 2]. ScÅtter – The SIBYLS Beamline [Internet]. Available from: https://bl1231.als.lbl.gov/scatter/

38. Hansen S. 2000. Bayesian estimation of hyperparameters for indirect Fourier transformation in small-angle scattering. J Appl Crystallogr. 33:1415–21.

39. Hopkins JB, Gillilan RE, Skou S. 2017. BioXTAS RAW: Improvements to a free open-source program for small-angle X-ray scattering data reduction and analysis. J Appl Crystallogr. 50:1545–53.

40. Abraham MJ, Murtola T, Schulz R, Páll S, Smith JC, Hess B, Lindah E. 2015. GROMACS: High performance molecular simulations through multi-level parallelism from laptops to supercomputers. SoftwareX. 1–2:19–25.

41. Bonomi M, Bussi G, Camilloni C, Tribello GA, Banáš P, Barducci A, et al. 2019. Promoting transparency and reproducibility in enhanced molecular simulations. Nat Methods. 16:670–3.

42. Abascal JLF, Vega C. 2005. A general purpose model for the condensed phases of water: TIP4P/2005. J Chem Phys. 123.

43. Piana S, Donchev AG, Robustelli P, Shaw DE. 2015. Water dispersion interactions strongly influence simulated structural properties of disordered protein states. Journal of Physical Chemistry B. 119:5113–23.

44. Jorgensen WL, Chandrasekhar J, Madura JD, Impey RW, Klein ML. 1983. Comparison of simple potential functions for simulating liquid water. J Chem Phys. 79:926.

45. Schrödinger, L., & DeLano W. 2020. PyMOL [Internet]. Available from: http://www.pymol.org/pymol

46. Hopkins CW, Le Grand S, Walker RC, Roitberg AE. 2015. Long-Time-Step Molecular Dynamics through Hydrogen Mass Repartitioning. J Chem Theory Comput. 11:1864–74.

47. Bussi G, Donadio D, Parrinello M. 2007. Canonical sampling through velocity rescaling. J Chem Phys. 126:014101.

48. Parrinello M, Rahman A. 1998. Polymorphic transitions in single crystals: A new molecular dynamics method. J Appl Phys. 52:7182.

49. Hess B, Bekker H, Berendsen HJC, Fraaije JGEM. 1997. LINCS: A Linear Constraint Solver for Molecular Simulations. J Comput Chem. 18:14631472.

50. Essmann U, Perera L, Berkowitz ML, Darden T, Lee H, Pedersen LG. 1998. A smooth particle mesh Ewald method. J Chem Phys. 103:8577.

51. Bussi G, Tribello GA. 2019. Analyzing and Biasing Simulations with PLUMED. Methods in molecular biology. 2022:529–78.

52. McGibbon RT, Beauchamp KA, Harrigan MP, Klein C, Swails JM, Hernández CX, et al. 2015. MDTraj: A Modern Open Library for the Analysis of Molecular Dynamics Trajectories. Biophys J. 109:1528.

53. Kohlhoff KJ, Robustelli P, Cavalli A, Salvatella X, Vendruscolo M. 2009. Fast and accurate predictions of protein NMR chemical shifts from interatomic distances. J Am Chem Soc. 131:13894–5.

54. Grudinin S, Garkavenko M, Kazennov A. 2017. Pepsi-SAXS: an adaptive method for rapid and accurate computation of small-angle X-ray scattering profiles. Acta Crystallogr D Struct Biol. 73:449–64.

55. Rauscher S, Gapsys V, Gajda MJ, Zweckstetter M, De Groot BL, Grubmüller H. 2015. Structural Ensembles of Intrinsically Disordered Proteins Depend Strongly on Force Field: A Comparison to Experiment. J Chem Theory Comput. 11:5513–24.

56. Piana S, Robustelli P, Tan D, Chen S, Shaw DE. 2020. Development of a Force Field for the Simulation of Single-Chain Proteins and Protein-Protein Complexes. J Chem Theory Comput. 16:2494–507.

57. Sormanni P, Camilloni C, Fariselli P, Vendruscolo M. 2015. The s2D Method: Simultaneous Sequence-Based Prediction of the Statistical Populations of Ordered and Disordered Regions in Proteins. J Mol Biol. 427:982–96.

58. Jumper J, Evans R, Pritzel A, Green T, Figurnov M, Ronneberger O, et al. 2021. Highly accurate protein structure prediction with AlphaFold. Nature. 596:583–9.

59. Wilson CJ, Choy WY, Karttunen M. 2022. AlphaFold2: A Role for Disordered Protein/Region Prediction? Int J Mol Sci. 23.

60. Eastman P, Swails J, Chodera JD, McGibbon RT, Zhao Y, Beauchamp KA, et al. 2017. OpenMM 7: Rapid development of high performance algorithms for molecular dynamics. PLoS Comput Biol. 13:e1005659.

61. Tesei G, Lindorff-Larsen K. 2023. Improved predictions of phase behaviour of intrinsically disordered proteins by tuning the interaction range. Open Research Europe. 2:94.

62. Tesei G, Trolle AI, Jonsson N, Betz J, Knudsen FE, Pesce F, Johansson KE, Lindorff-Larsen K. 2024. Conformational ensembles of the human intrinsically disordered proteome. Nature. 626:897–904.

63. Lotthammer JM, Ginell GM, Griffith D, Emenecker RJ, Holehouse AS. 2024. Direct prediction of intrinsically disordered protein conformational properties from sequences. Nat Methods. 21:465–76.

64. Baxter NJ, Williamson MP. 1997. Temperature dependence of 1H chemical shifts in proteins. Journal of Biomolecular NMR 1997 9:4. 9:359–69.

65. Grzesiek S, Anglister J, Bax A. 1993. Correlation of Backbone Amide and Aliphatic Side-Chain Resonances in 13C/15N-Enriched Proteins by Isotropic Mixing of 13C Magnetization. J Magn Reson B. 101:114–9.

66. Shen Y, Bax A. 2010. Prediction of Xaa-Pro peptide bond conformation from sequence and chemical shifts. J Biomol NMR. 46:199–204.

67. Hansen DF, Led JJ. 2003. Implications of using approximate Bloch–McConnell equations in NMR analyses of chemically exchanging systems: application to the electron self-exchange of plastocyanin. Journal of Magnetic Resonance. 163:215–27.

68. Hazra MK, Gilron Y, Levy Y. 2023. Not Only Expansion: Proline Content and Density Also Induce Disordered Protein Conformation Compaction. J Mol Biol. 435:168196.

69. Kay LE, Torchia DA, Bax A. 1989. Backbone Dynamics of Proteins As Studied by 15N Inverse Detected Heteronuclear NMR Spectroscopy: Application to Staphylococcal Nuclease. Biochemistry. 28:8972–9.

70. Korzhnev DM, Skrynnikov NR, Millet O, Torchia DA, Kay LE. 2002. An NMR experiment for the accurate measurement of heteronuclear spin-lock relaxation rates. J Am Chem Soc. 124:10743–53.

71. Abyzov A, Salvi N, Schneider R, Maurin D, Ruigrok RWH, Jensen MR, Blackledge M. 2016. Identification of Dynamic Modes in an Intrinsically Disordered Protein Using Temperature-Dependent NMR Relaxation. J Am Chem Soc. 138:6240–51.

