## Supplementary material for "An integrative characterisation of proline *cis* and *trans* conformers in a disordered peptide": orf6_str_si

### SUPPLEMENTARY METHODS

#### NMR assignment experiments

The 2D $^1$H-$^1$H total correlation spectroscopy (TOCSY) and 2D $^1$H-$^{13}$C heteronuclear single quantum coherence (HSQC) spectra were measured at a static magnetic field strength of 14.1 T (600 MHz) on the unlabelled N-acetylated (NAc-ORF6$_{CTR}$) to assign the $^1$H$^\alpha$, $^1$H$^\beta$, $^{13}$C$^\alpha$, and $^{13}$C$^\beta$ chemical shifts. These were used for comparison with the chemical shifts derived from the all-atom explicit solvent metadynamics simulations. The 2D $^1$H-$^1$H TOCSY spectra were acquired using the standard dipsi2esgpph Bruker pulse sequence (1,2), with spectral widths of 14 ppm (direct dimension) and 12 ppm (indirect dimension), and 1,024 x 512 complex points, respectively. 32 scans were recorded, with an inter-scan delay of 1.5 s. The RF-carrier was set to the position of the water peak ($^1$H; 4.77 ppm) in both dimensions. The TOCSY mixing was applied with a field of 10 kHz for 80 ms. This experiment was repeated on the ORF6$_{CTR}$ sample using the same setup. For the 2D $^1$H-$^{13}$C spectra, the standard hsqcgpph Bruker sequence was employed, with 192 scans, spectral widths of 14 ppm ($^1$H) and 80 ppm for ($^{13}$C), and 512 x 180 complex points, respectively. The recycle delay was set to 1.5 s, with carriers set to the position of the water peak ($^1$H; 4.77 ppm) and 35 ppm relative to TMS ($^{13}$C). These experiments were measured at temperatures ranging from 5˚C to 40˚C, increasing by 5˚C increments, and at 37˚C.

The 2D $^1$H-$^{15}$N HSQC spectra were recorded on the uniformly $^{15}$N-labelled ORF6$_{CTR}$ at a static magnetic field strength of 14.1 T and at temperatures ranging from 5˚C to 40˚C, increasing by 5˚C increments. Additionally, measurements were taken at 2.5˚C and 37˚C. The 2D $^1$H-$^{15}$N HSQC spectra were acquired using the hsqcetf3gpsi2 Bruker pulse sequence (3–5). Spectral widths were set to 16 ppm ($^1$H) and 20.5 ppm ($^{15}$N), with 1,536 x 128 complex points acquired, respectively. Carriers were set to the position of the water peak ($^1$H; 4.77 ppm) and 121.5 ppm ($^{15}$N), with 4 scans and a recycle delay of 1 s.

Uniformly $^{13}$C,$^{15}$N-labelled and $^{15}$N-labelled samples of ORF6$_{CTR}$ were used for backbone and side-chain chemical shift assignments. These assignments correspond to both the *trans*-P57 and *cis*-P57 conformations at a static magnetic field strength of 14.1 T and at 15°C. Standard double-resonance and triple-resonance experiments were employed for backbone assignment, including; the CON, HNCO (6,7), HN(CA)CO (8), and HNCACB (9) experiments. This was supplemented with the $^{15}$N-edited nuclear Overhauser effect spectroscopy HSQC ($^{15}$N-NOESY-HSQC) (10,11) and $^{15}$N-TOCSY-HSQC (12) experiments to characterise intra- and inter-residue information. Side-chain assignments were obtained from CC-TOCSY-type experiments, which included the CC(CO)NH (13) and the H(C)CH-TOCSY (14,15) experiments.

The 2D CON spectra were acquired using the c_con_iasq Bruker pulse sequence (16,17). Spectral widths were set to 40 ppm in both dimensions, with carriers set to 173 ppm relative to TMS ($^{13}$C) and 121.5 ppm ($^{15}$N), and 512 x 128 complex points, respectively. 32 scans and a recycle delay of 2 s were applied. The 3D HNCO spectra were acquired using the hncogp3d Bruker pulse sequence (5,18) and the 3D HN(CA)CO spectra were acquired using the hncacogpwg3d Bruker pulse sequence (8). Both spectra were recorded with 16 scans, 1,024 x 29 x 80 complex points ($^1$H, $^{15}$N, $^{13}$C), and a spectral width of 14 x 20.5 x 7 ppm, respectively. Carriers were set to the position of the water peak ($^1$H; 4.77 ppm), 121.5 ppm ($^{15}$N) and 173 ppm relative to TMS ($^{13}$C), with a recycle delay of 1 s. The 3D HNCACB spectra were acquired using the hncacbgp3d Bruker pulse sequence (9,19). Spectra were recorded with 16 scans, 1,024 x 30 x 72 complex points ($^1$H, $^{15}$N, $^{13}$C), and a spectral width of 14 x 20.5 x 60.2 ppm, respectively. Carriers were set to the water peak ($^1$H; 4.77 ppm), 121.5 ppm ($^{15}$N), and 43 ppm ($^{13}$C), with a recycle delay of 1 s.

The $^{15}$N-NOESY-HSQC spectra were acquired using the noesyhsqcfpf3gpsi3d Bruker pulse sequence (3,5). Spectra were recorded with 8 scans, 2,048 x 24 x 96 complex points ($^1$H, $^{15}$N, $^1$H), and a spectral width of 16 x 20.5 x 16 ppm, respectively. Carriers were set to the water peak ($^1$H; 4.77 ppm) and 121.5 ppm ($^{15}$N). A recycle delay of 1.5 s and a mixing time of 0.25 s were used. The $^{15}N$-TOCSY-HSQC spectra were acquired using a pulse sequence based on the original Bruker dipsihsqcf3gpsi3d with a flip-flop spectroscopy (FLOPSY)-16 scheme (3–5,12,20). Spectra were acquired with 16 scans, 1,024 x 32 x 96 complex points ($^{1}H$, $^{15}N$, $^{1}H$), and a spectral width of 16 x 20.5 x 16 ppm, respectively. Carriers were set to the water peak ($^{1}H$; 4.77 ppm) and 121.5 ppm ($^{15}N$), with a recycle delay of 1 s. A TOCSY mixing time of 0.1 s was used and an 8 kHz TOCSY spin lock was applied.

The CC(CO)NH spectra were acquired using a pulse sequence based on the original Bruker ccconhgp3d.2 with a FLOPSY-16 scheme (20). Spectra were recorded with 16 scans, 1,024 x 30 x 84 complex points ($^{1}H$, $^{15}N$, $^{13}C$), and a spectral width of 14 x 20.5 x 71 ppm, respectively. Carriers were set to the water peak ($^{1}H$; 4.77 ppm), 121.5 ppm ($^{15}N$) and 43 ppm ($^{13}C$). A recycle delay of 1.5 s, along with a TOCSY mixing time of 18 ms were used. The H(C)CH-TOCSY spectra were acquired using a pulse sequence based on the original Bruker hcchdigp3d with a FLOPSY-16 scheme (14,20). Spectra were recorded with 32 scans, 1,024 x 24 x 112 complex points ($^{1}H$, $^{1}H$, $^{13}C$), and a spectral width of 15 x 12 x 69 ppm, respectively. Carriers were set to the water peak ($^{1}H$; 4.77 ppm) and 43 ppm ($^{13}C$). A recycle delay of 1.5 s was used, along with a TOCSY mixing time of 18 ms, and a $^{13}C$ spin-lock pulse applied at 8 kHz.

**NMR relaxation experiments**

$^{15}N$-labelled ORF6$_{CTR}$ samples were used to measure the backbone longitudinal ($R_1$) and rotating frame ($R_{1\rho}$) relaxation rates, as well as the heteronuclear $\{^{1}H\}$-$^{15}N$ steady-state NOEs (hetNOEs), at two static magnetic field strengths (14.1 T and 18.8 T) and at 15˚C.

Both $R_1$ and $R_{1\rho}$ were measured using established proton-detected pulse sequences based on a gradient-selected, sensitivity-enhanced, refocused $^{15}N$ sequences (21,22). Both spectra were recorded with 2,048 x 160 complex points and spectral widths as used before in the 2D $^{1}H$-$^{15}N$ HSQC experiment. Gradient pulses were used to suppress the water signal and a long recycle delay of 3 s was employed. In the $R_1$ sequence, N-H cross-correlated relaxation pathways were suppressed by hard 180˚ pulses every 20 ms during the relaxation delay (23,24). The $R_1$ $^{15}N$-$^{1}H$ planes were recorded with 8 relaxation delays ranging from 20 ms to 700 ms. In the $R_{1\rho}$ sequence, cross-correlated relaxation was suppressed by hard 180˚ pulses during the $^{15}N$ spin-lock (25). Magnetisation was explicitly aligned with the spin-lock field (26). N-H planes were recorded using 8 relaxation delays ranging from 2 ms to 140 ms, with a $^{15}N$ spin-lock field strength of 2 kHz.

The $\{^{1}H\}$-$^{15}N$ hetNOEs were recorded using a pseudo-3D experiment, with and without proton saturation (21,22,27). Amide proton magnetisation saturation was accomplished by using a 5 s train of high-power 120° pulses applied at 5 ms intervals. The reference and saturated spectra were alternately recorded. To ensure complete recovery of the initial magnetisation at the start of each increment of the reference experiment, a long recycle delay of 15 s was applied (28). The spectral widths were 16 ppm ($^{1}H$) and 20.5 ppm ($^{15}N$), and 2,048 x 160 ($^{1}H$, $^{15}N$) complex points were recorded at a static magnetic field strength of 14.1 T.

Exchange-free $^{15}N$ transverse relaxation ($R_{dd}$) rates were measured using pulse schemes for the four $^{1}H$-$^{15}N$ relaxation rates $R_{1\rho}(2H_zN'_z)$, $R_{1\rho}(2H'_zN_z)$, $R_{1\rho}{}^2(2H'_zN'_z)$, and $R_1(2H_zN_z)$ at a static magnetic field strength of 14.1 T and at 15˚C (29). A total of 8 relaxation delays between 2 ms and 26 ms were used for all experiments (same delays used for each rate measurement). A 10 kHz $^{1}H$ spin lock and a 2 kHz $^{15}N$ spin lock were applied. Spectral widths were the same as those used before in $\{^{1}H\}$-$^{15}N$ hetNOE experiments, with 1,536 x 640 ($^{1}H$, $^{15}N$) complex points.

**NMR diffusion experiments**

$^{15}N$-labelled ORF6$_{CTR}$ samples were used to measure the pseudo-3D diffusion pulsed field-gradient spin echo (PFGSE) $^{1}H$-$^{15}N$-Diffusion Ordered Spectroscopy-HSQC ($^{1}H$-$^{15}N$-DOSY-HSQC) experiment with a bipolar gradient (30) at a static magnetic field strength of 22.3 T and at 15˚C. Six experiments with gradient strengths of

0.963, 8.667, 16.37, 24.07, 31.78, and 39.48 G/cm were acquired for each data set, with the gradient strengths augmented linearly through the acquisition and all other delays and pulses held constant. Gradient pulses ($\delta$) were applied for 3 ms and a diffusion delay ($\Delta$) of 200 ms. 80 scans were acquired per gradient experiment with 1,536 x 176 complex points ($^1$H, $^{15}$N), using the same spectral widths and carriers as used before for the 2D $^1$H-$^{15}$N HSQC experiment.

**NMR analysis**

The transverse relaxation rate-constants ($R_2$) were calculated using the following equation:

$$R_{1\rho} = R_1 \cos^2\theta + R_2 \sin^2\theta \tag{S1}$$

Where $\theta$ is the tip-angle of the magnetisation-vector:

$$\theta = \arctan\left(\frac{\nu_1}{\Delta\nu}\right) \tag{S2}$$

and $\nu_1$ is the field strength of the spin-lock (Hz) and $\Delta\nu$ is the offset relative to the frequency of the transmitter frequency (Hz).

Diffusion coefficients of the *cis*-P57 and *trans*-P57 peaks were calculated by fitting the signal decay of residues Q56, M58, and E59 against the Stejskal-Tanner equation (31):

$$\frac{I}{I_0} = e^{-D\gamma_{\mathrm{H}}^2 g^2 \delta^2\left(\Delta-\frac{\delta}{3}-\frac{\tau}{2}\right)} \tag{S3}$$

where $I$ is the observed intensity, $I_0$ is the intensity of the unattenuated signal, $D$ is the diffusion coefficient, and $\gamma_{\mathrm{H}}$ is the gyromagnetic ration of $^1$H. Residue E55 was not included in the analysis due to low signal intensity.

**Metadynamics setup**

The following 9 collective variables (CVs) were used to enhance conformational sampling of the NAc-ORF6$_{\mathrm{CTR}}$ with Gaussian widths for each CV shown in Table S1 (32,33):

1.) Total α-helix content, where the ALPHARMSD keyword in PLUMED is used to quantify this CV (34). In proteins, any chain of six contiguous residues can form an α-helix. This CV initially generates the set of all potential regions comprising six consecutive residues within the system. The root mean square deviation (RMSD) distance between the configuration where the residues are located and an idealised helix structure is then calculated. The sum of functions of the RMSD distances are used to compute this CV as follows:

$$S_{\alpha\mathrm{helix}} = \Sigma_i \frac{1-\left(\frac{\mathrm{RMSD}_i}{\mathrm{RMSD}_0}\right)^8}{1-\left(\frac{\mathrm{RMSD}_i}{\mathrm{RMSD}_0}\right)^{12}} \tag{S4}$$

where the sum extends over all potential α-helix segments, $i$, and $\mathrm{RMSD}_0$ = 0.08 nm.

2.) The sum of the combined parallel and anti-parallel β-sheet content, where the total parallel β-sheet content and total anti-parallel β-sheet content are calculated using the PARABETARMSD and ANTIBETARMSD keywords, respectively, in PLUMED. In a protein chain, two segments containing three contiguous residues can form an anti-parallel or parallel β-sheet if they are separated by a minimum of two or three residues, respectively, to accommodate a turn. This CV initially generates the set of all possible six residue sections that could form a parallel or anti-parallel β-sheet. It then calculates the RMSD distances between the configuration where the residues are located and the idealised parallel or anti-parallel β-sheet structure. The total parallel or anti-parallel β-sheet content is calculated using the equivalent Eq. S4 above, where the sum extends over all potential parallel and anti-parallel β-sheets. The CV is finally computed by summing the combined parallel and anti-parallel β-sheet contents (34).

3.) The $R_g$ computed on the $C^\alpha$ carbons, using the GYRATION keyword in PLUMED. This CV is defined as

$$S_{R_g} = \left( \frac{\sum_{i \in C_\alpha} m_i |r_i - r_{COM}|^2}{\sum_{i \in C_\alpha}^{n} m_i} \right)^{\frac{1}{2}} \tag{S5}$$

where $m_i$ and $r_i$ are the mass and position of the $C^\alpha$ atom of the $i^{th}$ residue, respectively.

$\boldsymbol{r}_{COM}$ is the coordinate of the centre of mass (COM) of the protein defined as:

$$r_{COM} = \frac{\Sigma_{i \varepsilon C^\alpha} m_i \boldsymbol{r}_i}{\Sigma_{i \varepsilon C^\alpha} m_i} \tag{S6}$$

4.) The number of salt bridges. This CV is computed as the number of contacts within a 0.6 nm cut-off range between the atoms of $NH_3^+$ groups of lysine or the $-C(NH_2)_2^+$ of arginine and the $-COO^-$ atom groups of aspartate or glutamate. This CV is computed using the COORDINATION key word in PLUMED (33). The following switching function was used:

$$S_{\text{Salt bridges}} = \sum_{i \in A} \sum_{j \in B} \frac{1 - \left(\frac{d_{ij}}{d_0}\right)^6}{1 - \left(\frac{d_{ij}}{d_0}\right)^{12}} \tag{S7}$$

where $d_0$ = 0.6 nm, $d_{ij}$ is the distance between atoms $i$ and $j$, and A and B are the two groups of atoms between which contacts are calculated. For this example, group A contains the heavy atoms from arginine and lysine residues and group B contains the heavy atoms from aspartate and glutamate residues. Self-interactions are excluded from the calculation.

5.) The end-to-end distance of the NAc-ORF6$_{CTR}$. This CV is computed on the $C^\alpha$ carbon of the first and last residue using the DISTANCES keyword in PLUMED, which calculates the distance between a pair of atoms.

6.) The correlation between consecutive $\psi$ torsion angles (33). This CV is computed using the DIHCOR keyword in PLUMED applying the following equation for every residue, *i*:

$$s_{\text{Dihedral correlation}} = \frac{1}{2}\sum_{i}[1 + \cos(\psi_i - \psi_{i+1}) \quad \text{(S8)}$$

7.) The improper dihedral angle $\zeta$ for proline. This CV uses the TORSION keyword in PLUMED to compute the torsion between the following four atoms: $C^{\alpha}_{i-1}$, $O_{i-1}$, $C^{\delta}_{i}$, $C^{\alpha}_{i}$, where *i* is the proline residue (35,36).

8.) The $\psi$ torsional angle for proline. This CV uses the TORSION keyword in PLUMED to compute the torsion between the following four atoms: $N_i$, $C^{\alpha}_{i}$, $C'_i$, $N_{i+1}$, where $i$ is the proline residue (35,36).

9.) The number of contacts between hydrophobic residues (33). Hydrophobic residues isoleucine, leucine, methionine, proline, and tyrosine are used in the CV, which calculates the number of inter-$C^{\beta}$ atom distances less than 0.6 nm. To compute the CV, the COORDINATION keyword in PLUMED is applied and uses the switching function as defined by Eq. S7 above, except A and B are the same group. Self-interactions are not included in the calculation.

**Convergence model**

The following equation was used to fit a bi-exponential curve to the FES blocking analysis for each CV:

$$\text{Error (FES)} = A * \left(1 - B * e^{(-k1*t)} - C * e^{(-k2*t)}\right) \quad \text{(S9)}$$

The constant term, *A*, determined from the model, was used to approximate the FES standard-error at infinite time for each CV.

**Structural ensemble analysis**

By using the weights $w(\mathbf{s}_i)$ obtained from the bias potential $V_{\text{PB}}(\mathbf{s}_i)$ at the end of the metadynamics simulations, observables were calculated as ensemble averages using the following equation:

$$w(\mathbf{s}_i) = \frac{e^{\frac{V_{\text{PB}}(\mathbf{s}_i)}{k_{\text{B}}T}}}{\sum_j^N e^{\frac{V_{\text{PB}}(\mathbf{s}_j)}{k_{\text{B}}T}}} \quad \text{(S10)}$$

where $\mathbf{s}_i$ is the value of all CVs at time $i$, $N$ is the total number of time steps, and $k_{\text{B}}T$ is 2.5774 kJ·mol$^{-1}$ (the product of the Boltzmann constant ($k_{\text{B}}$) and $T$, when $T$ = 310.15 K) (33,37).

### Supplementary Tables

**Table S1. Gaussian widths and grid sizes for the 9 CVs in the NAc-ORF6$_{CTR}$ metadynamics simulations for both a03ws simulations (run 1 and run 2), the a99SB-*disp* simulation, and the C36m simulation.**

| | Gaussian width (in CV units) | | | | |
|---|---|---|---|---|---|
| **CV** | **a03ws** | **a99SB-*disp*** | **C36m** | **GRID MAX** | **GRID MIN** |
| **1. α-helix** | 0.73 | 0.58 | 0.23 | 100 | -1 |
| **2. β-sheet** | 0.12 | 0.13 | 0.24 | 100 | -1 |
| **3. $R_g$** | 0.10 | 0.11 | 0.12 | 20 | -1 |
| **4. Salt bridges** | 2.47 | 2.61 | 4.01 | 400 | -1 |
| **5. End-to-end distance** | 0.48 | 0.54 | 0.63 | 10 | -1 |
| **6. Dihedral correlation** | 1.03 | 1.00 | 1.06 | 50 | -1 |
| **7. $\zeta$ angle** | 0.93 | 1.41 | 1.11 | π | -π |
| **8. $\psi$ angle** | 0.85 | 0.56 | 0.57 | π | -π |
| **9. Hydrophobic contacts** | 0.07 | 0.08 | 0.06 | 10 | -1 |

The upper and lower bounds of the grid are also provided. The upper and lower bounds for the grid are the maximum and minimum values that the CV can take, e.g., for the proline $\psi$ torsional angle for CV No. 8., the maximum value of the angle is π and the minimum value is -π.

**Table S2. Absolute differences for each residue in the ORF6$_{CTR}$ and NAc-ORF6$_{CTR}$ $^{1}H^{N}$ and $^{1}H^{\alpha}$ chemical shifts in ppm.**

| Residue | $\lvert\delta_H{}^{\alpha}\rvert$ (ppm) | $\lvert\Delta\delta_H{}^{N}\rvert$ (ppm) |
|---:|---:|---:|
| **S43** | 0.030 | 0.149 |
| **L44** | 0.018 | 0.092 |
| **T45** | 0.009 | 0.005 |
| **E46** | 0.002 | 0.008 |
| **N47** | 0.008 | 0.000 |
| **K48** | 0.003 | 0.005 |
| **Y49** | 0.002 | 0.001 |
| **S50** | 0.009 | 0.006 |
| **Q51** | 0.032 | 0.002 |
| **L52** | 0.005 | 0.007 |
| **D53** | 0.008 | 0.006 |
| **E54** | 0.013 | 0.001 |
| **E55** | 0.010 | 0.007 |
| **Q56** | 0.004 | 0.002 |
| **P57** | 0.018 | N/A |
| **M58** | 0.009 | 0.004 |
| **E59** | 0.002 | 0.003 |
| **I60** | 0.022 | 0.009 |
| **D61** | 0.017 | 0.010 |

The RMSD between ORF6$_{CTR}$ and NAc-ORF6$_{CTR}$ is 0.010 and 0.029 for $^{1}H^{\alpha}$ and $^{1}H^{N}$, respectively, as given by Eq. S11 shown below.

$$\mathrm{RMSD}(\mathbf{v},\mathbf{w}) = \left(\frac{1}{N}\sum_{i=1}^{N}(v_i - w_i)^2\right)^{\frac{1}{2}} \tag{S11}$$

where **v** = $\mathrm{ORF6_{CTR}}$ and **w** = NAc-$\mathrm{ORF6_{CTR}}$ for the $^1\mathrm{H^N}$ or $^1\mathrm{H^{\alpha}}$ chemical shifts assigned for residues 43-61.

### SUPPLEMENTARY FIGURES

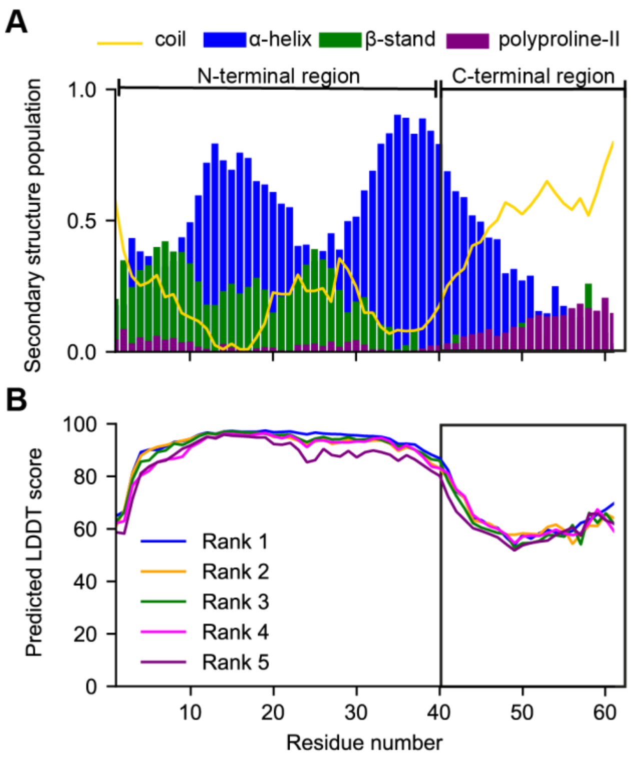


**Figure S1. The C-terminal region of ORF6 is predicted to be intrinsically disordered.** (*A*) s2D prediction showing the percentage populations of coil, α-helix, β-stand, and polyproline-II (38). (*B*) AlphaFold2 predicted local distance difference test (pLDDT) score (39). The pLDDT score is between 0 and 100 where pLDDT > 90 is a very high model confidence and pLDDT < 50 is a very low model confidence. The confidence for the $ORF6_{CTR}$ (residues 41-61) is very low indicating that this region is unstructured.

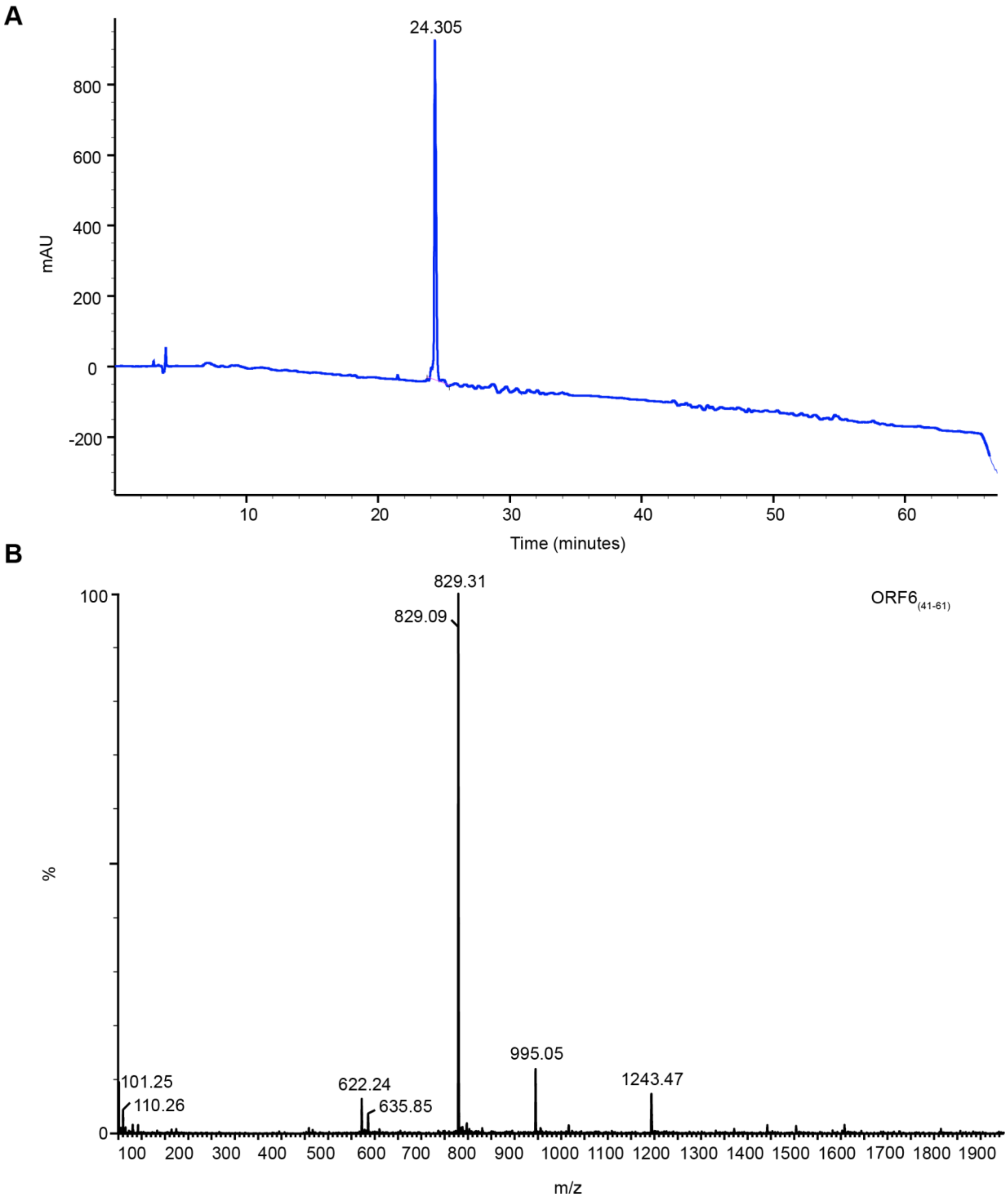


**Figure S2. ORF6$_{CTR}$ solid-phase peptide synthesis analysis.** (*A*) Analytical HPLC trace of purified ORF6$_{CTR}$ Analysis was performed using a gradient of Buffer B (0.1% (v/v) TFA in acetonitrile) in Buffer A (0.1% (v/v) TFA in water) (5-75% over 60 min) using a C8 RP-silica column from Dr. Maisch GmbH at 1 mL/min with detection at 214 nm. (*B*) ESI+ mass spectrum of purified ORF6$_{CTR}$ with annotations showing charge states. The spectrum was recorded with a Waters Acquity UPLC SQD LC-MS instrument equipped with a Hypersil GOLD C4, 5 µm particle, 50 mm x 2.1 mm column (Thermo Scientific).

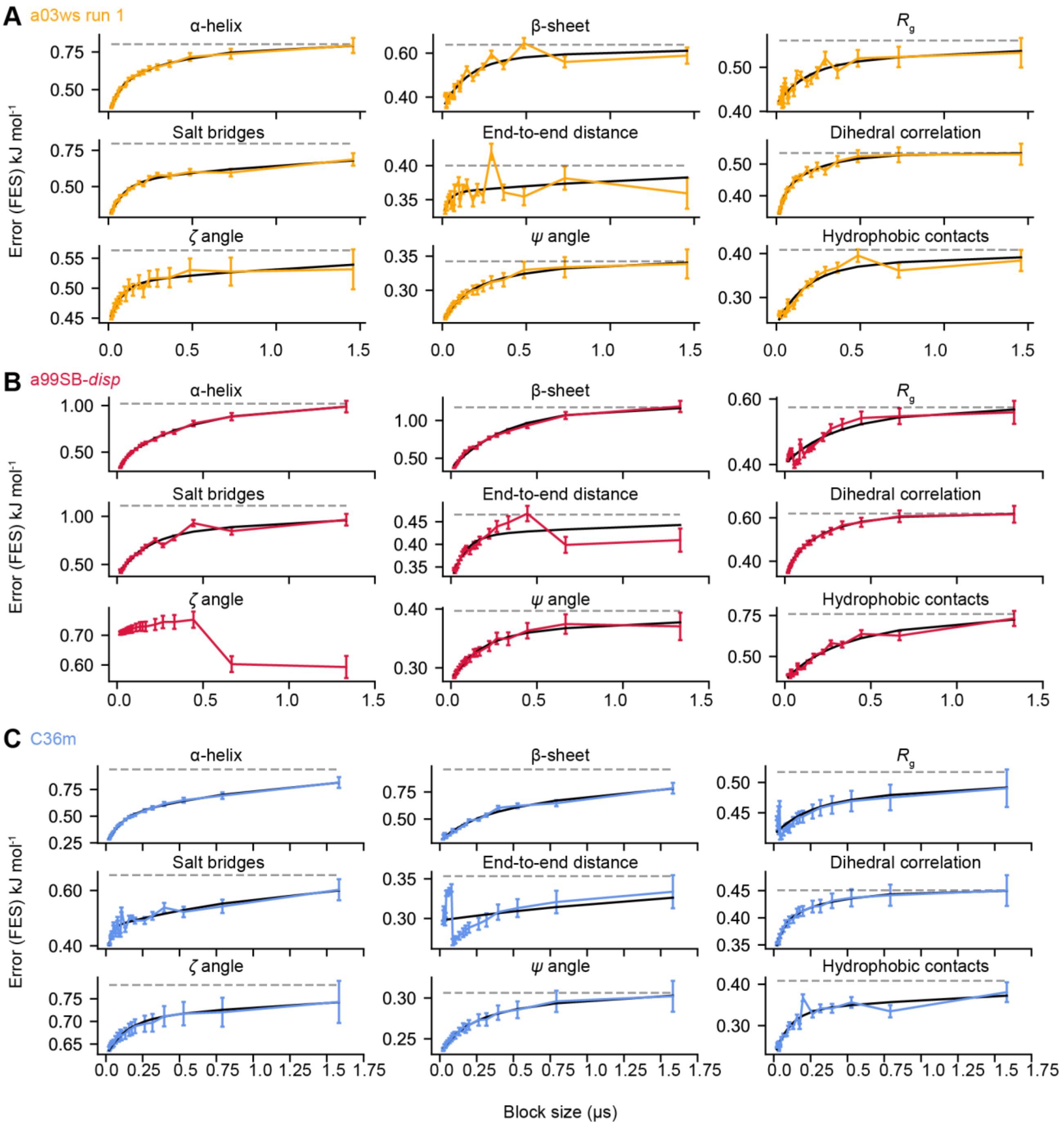


**Figure S3. Convergence analysis of NAc-ORF6$_{CTR}$ metadynamics simulations: a03ws run 1, a99SB-*disp,* and C36m ensembles.** The free energy surface (FES) standard-error as a function of block size was plotted for all 9 CVs in the (*A*) a03ws run 1 (orange), (*B*) a99SB-*disp* (red), and (*C*) C36m (blue) force fields (40,41). The modelled blocking analysis curve from Eq. S9 (black) and the FES standard-error at infinite time (grey dashed line) are shown. See below for a03ws run 2.

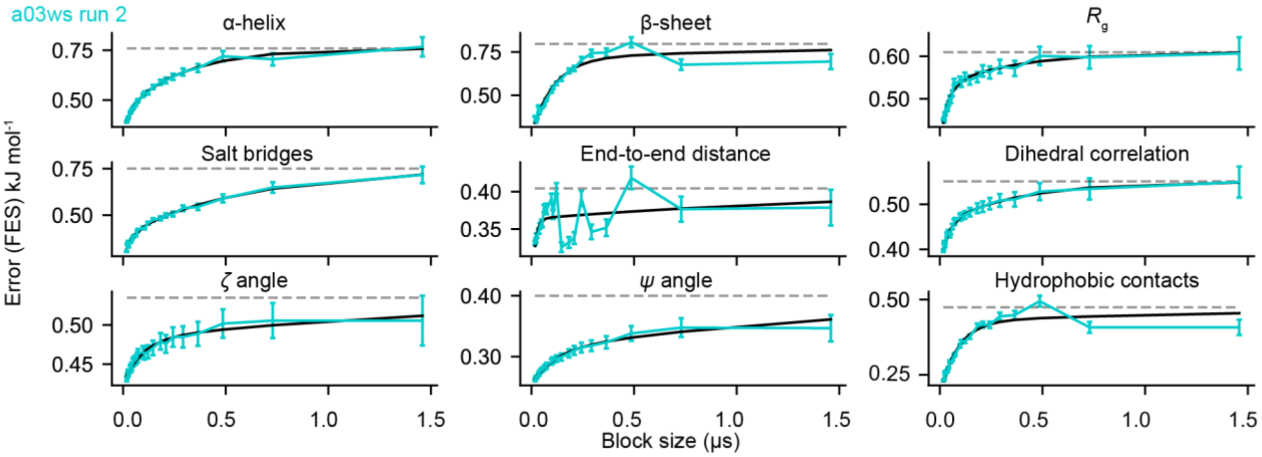


**Figure S4. Convergence analysis of NAc-ORF6$_{CTR}$ metadynamics simulations: a03ws run 2 ensemble.** The FES standard-error as a function of block size was plotted for all 9 CVs in the a03ws run 2 simulation (40,41). The modelled blocking analysis curve from Eq. S9 (black) and the FES standard-error at infinite time (grey dashed line) are shown.

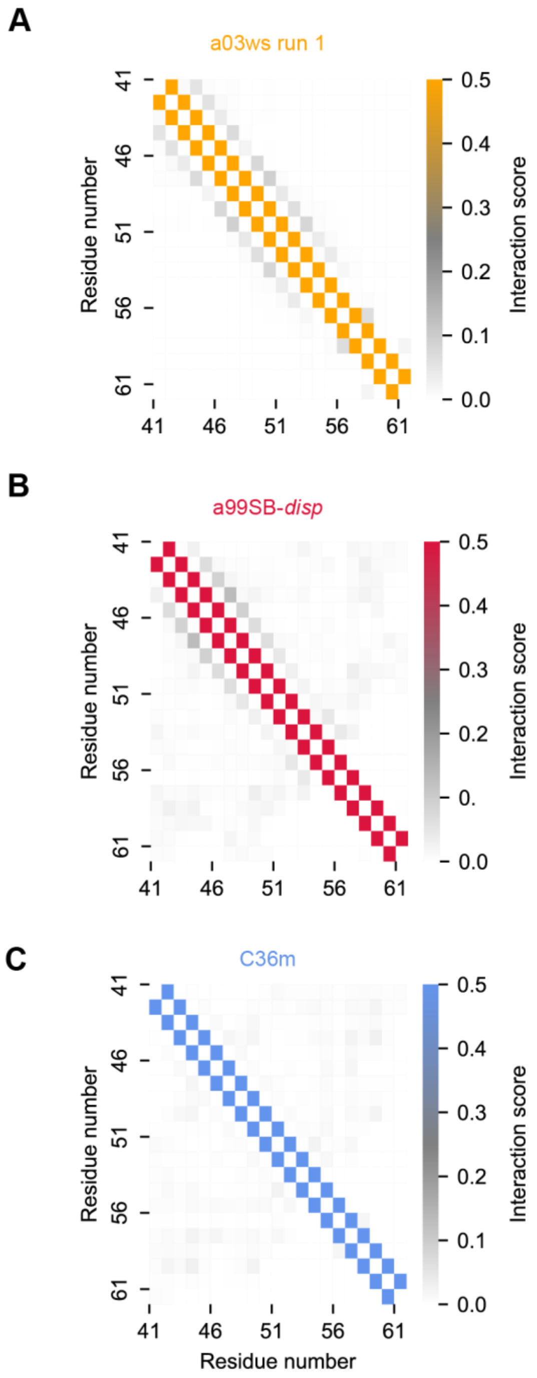


**Figure S5. C$^{\alpha}$ minimum distance inter-residue contact maps predict the NAc-ORF6$_{CTR}$ to be disordered in all three force fields (a03ws run 1, a99SB-*disp*, and C36m).** Contact maps are shown for (*A*) a03ws run 1 (orange), (*B*) a99SB-*disp* (red), and (*C*) C36m (blue) ensembles. The low probability of α-helix and β-sheet contacts indicates that all NAc-ORF6$_{CTR}$ conformational ensembles lack substantial secondary structure. The contact maps are for the metadynamics simulations before SAXS BME reweighting. a03ws run 2 is shown below.

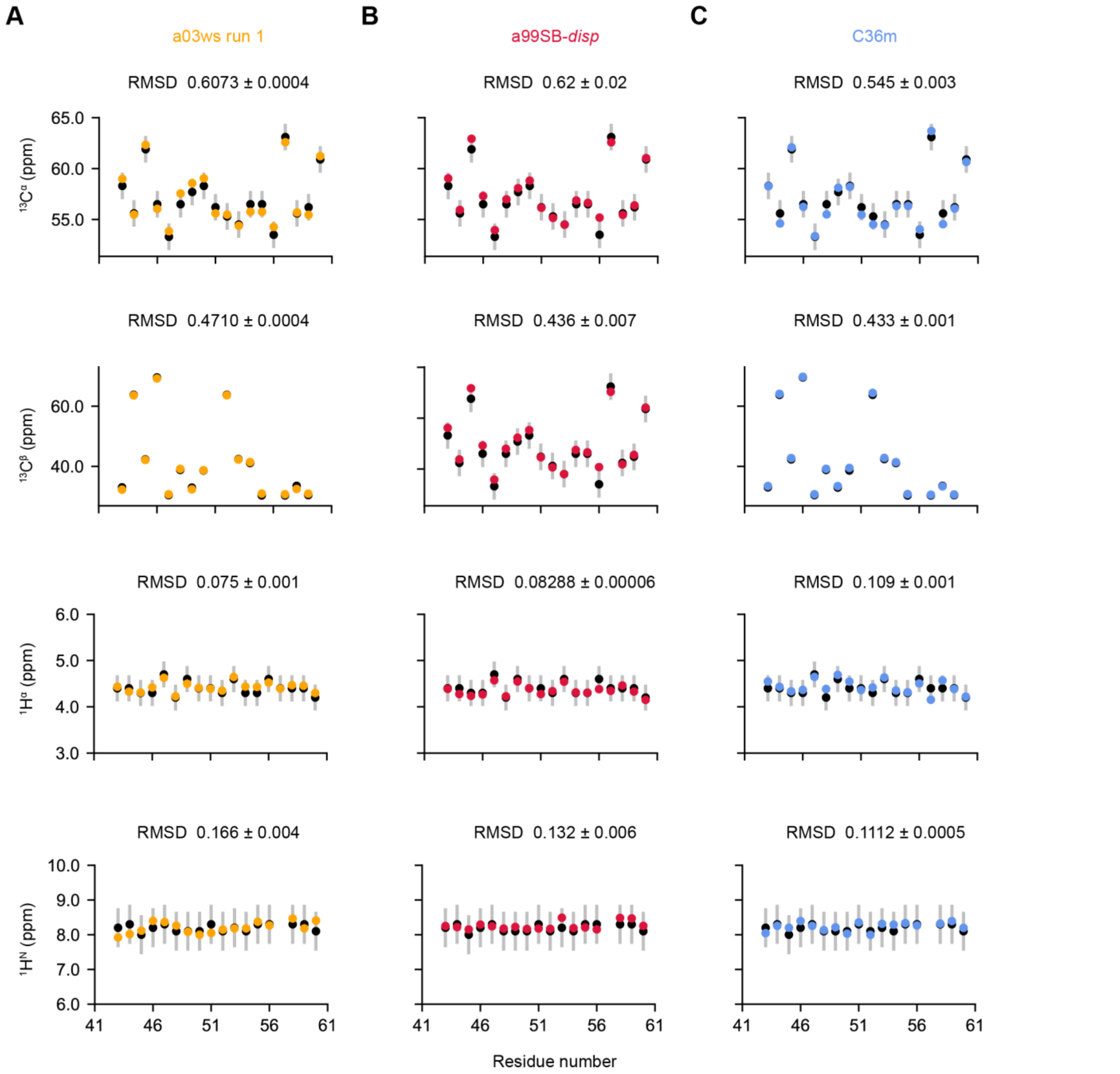


**Figure S6. Assessment of metadynamics simulations (before SAXS BME reweighting) consistency with NMR chemical shift data: a03ws run 1, a99SB-*disp,* and C36m ensembles.** Comparison of the ensemble-averaged predicted chemical shifts calculated using CamShift (42) for the (*A*) a03ws run 1 (orange), (*B*) a99SB-*disp* (red), and (*C*) C36m (blue) ensembles to experimental chemical shifts (black) using the $^{13}C^{\alpha}$, $^{13}C^{\beta}$, $^{1}H^{\alpha}$, and $^{1}H^{N}$ chemical shifts. The error in CamShift (silver) is shown. The standard deviation between the chemical shifts predicted in the first and second halves of the analysed trajectory were also plotted but are smaller than the data points. The RMSD and its corresponding error between experimental and chemical shifts predicted from the simulations are displayed for each force field and chemical shift.  a03ws run 2 is included below.

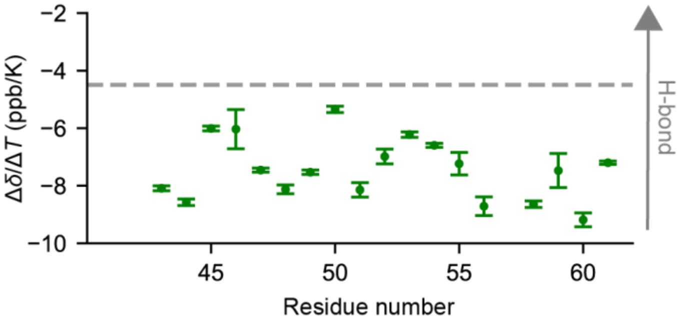


**Figure S7. NMR analysis indicates the amide protons in the NAc-ORF6$_{CTR}$ do not form stable hydrogen bonds.** Temperature coefficients for the *trans*-P57 conformation of 400 µM unlabelled NAc-ORF6$_{CTR}$ determined from $^1H^\alpha$-$^1H^N$ TOCSY spectra at 8 temperatures between 5-37˚C. A value more negative than -4.5 ppb/K (grey dashed line) indicates a lack of hydrogen bonding. Error bars represent the uncertainty associated with the least squares polynomial fit for each residue. Data was collected at pH 6.9 and at a static magnetic field strength of 14.1 T.

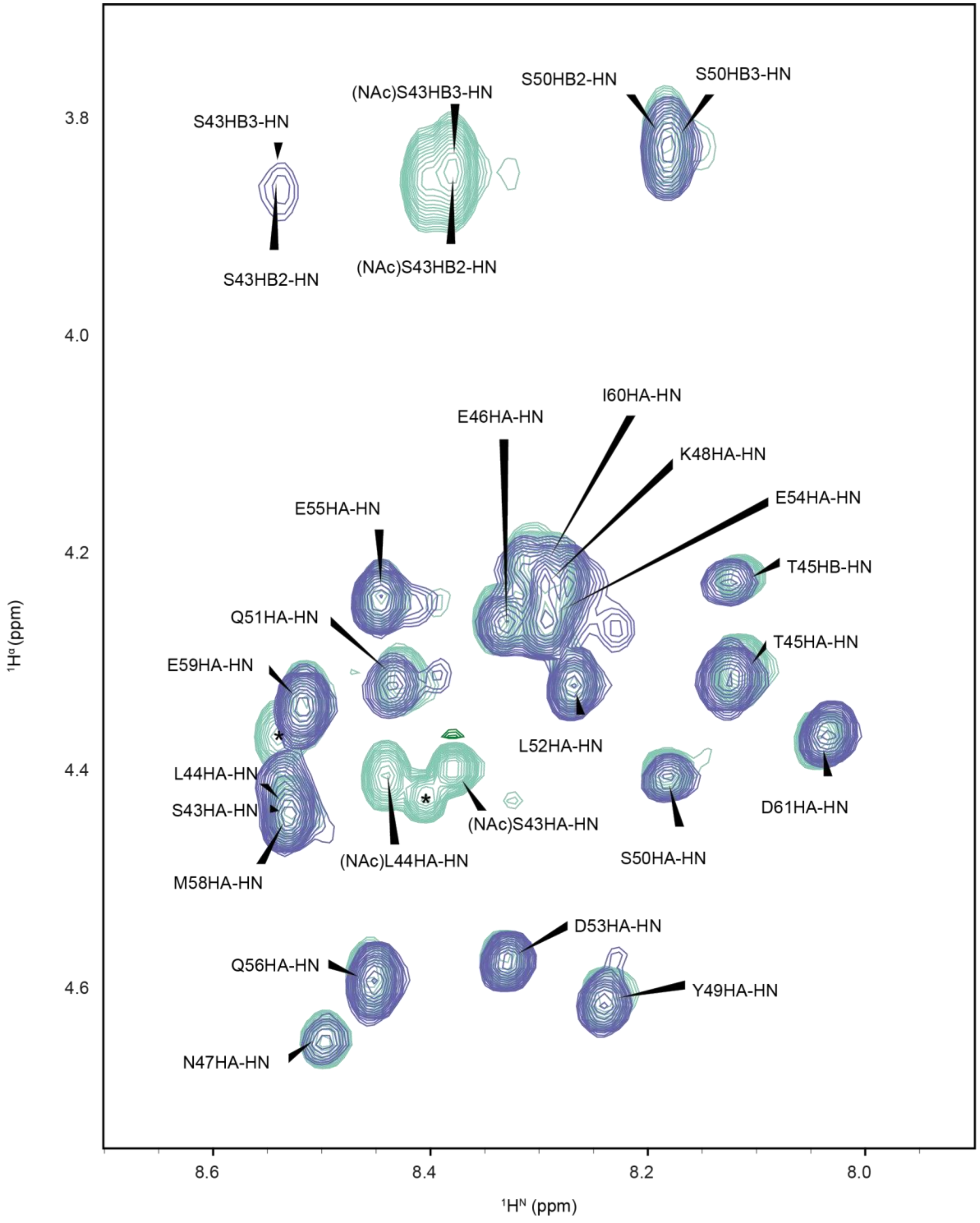


**Figure S8. ORF6$_{CTR}$ and NAc-ORF6$_{CTR}$ $^1$H$^\alpha$-$^1$H$^N$ TOCSY spectra.** $^1$H$^\alpha$-$^1$H$^N$ TOCSY for 300 µM $^{15}$N-labelled ORF6$_{CTR}$ (blue-purple) and 400 µM unlabelled NAc-ORF6$_{CTR}$ (green-cyan) at 15˚C. Narrow dispersion in the $^1$H$^N$ dimension indicates that ORF6$_{CTR}$ is disordered. N-acetylation only notably changes the chemical shifts for the residues closest to the N-terminus (S43 and L44). Spectra were acquired at a static magnetic field strength of 14.1 T and at pH 6.9. Unassigned peaks have been marked with an asterisk (*).

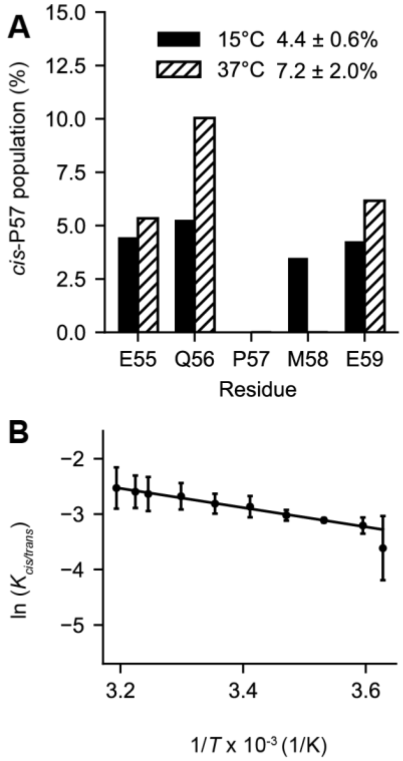


**Figure S9. ORF6$_{CTR}$ *cis*-P57 population and thermodynamics analysis using peak height.** (*A*) The *cis*-P57 populations at 15˚C (solid fill) and 37˚C (dashed fill) are shown for each well-resolved *cis*-P57 and *trans*-P57 peak in the 2D $^{1}H$-$^{15}N$ HSQC spectrum. The peak height was used to calculate the mean *cis*-P57 population and standard deviation across residues E55, Q56, and E59. (*B*) The Van't Hoff analysis of *cis/trans* P57 isomerisation determined from the peak height. The mean natural logarithm of the equilibrium constant for proline *cis*/*trans* isomerisation around the Q56-P57 bond as a function of temperature was calculated using residues E55, Q56, and E59. Error bars represent the standard deviation across the 3 residues used in the analysis. The Van't Hoff linear fit yielded a *cis*-P57 population of 8 ± 1% at 37˚C. Experiments were recorded at a static magnetic field strength of 14.1 T and at pH 6.9.

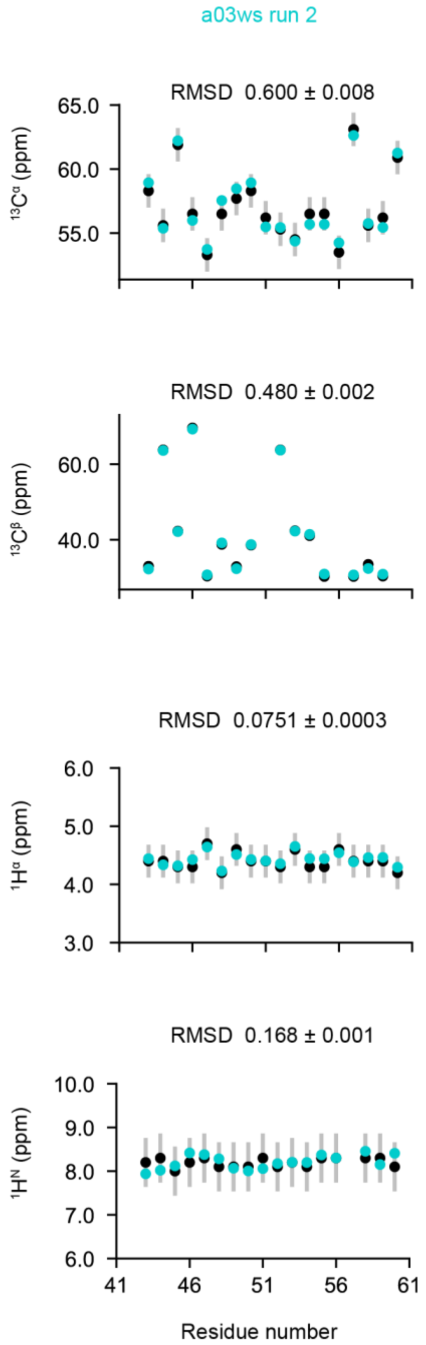


**Figure S10. Assessment of metadynamics simulations (before SAXS BME reweighting) consistency with NMR chemical shift data: a03ws run 2 ensemble.** Comparison of the ensemble-averaged predicted chemical shifts calculated using CamShift (42) for the a03ws run 2 (cyan) ensemble to experimental chemical shifts (black) using the $^{13}C^{\alpha}$, $^{13}C^{\beta}$, $^{1}H^{\alpha}$, and $^{1}H^{N}$ chemical shifts. The error in CamShift (silver) is shown. The standard deviation between the chemical shifts predicted in the first and second halves of the analysed trajectory were also plotted but are smaller than the data points. The RMSD and its corresponding error between experimental and chemical shifts predicted from the simulations are displayed for each force field and chemical shift.

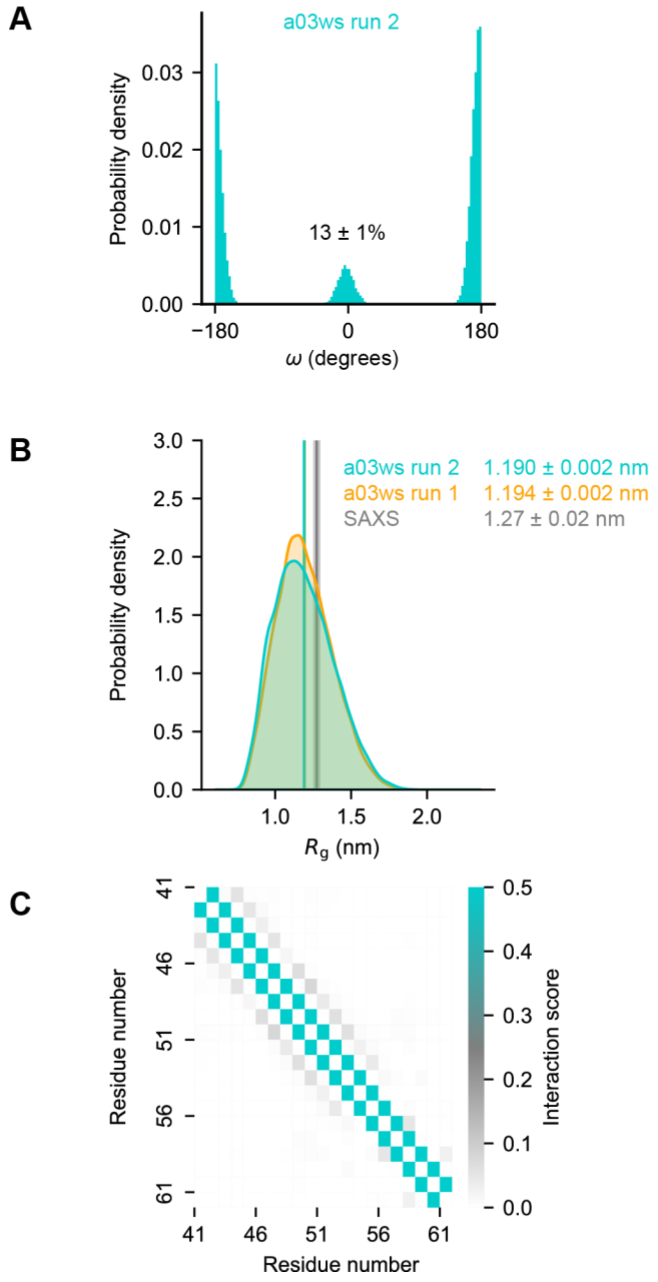


**Figure S11. Consistency of the a03ws run 2 NAc-ORF6$_{CTR}$ ensemble characterisations with the a03ws run 1 ensemble.** NAc-ORF6$_{CTR}$ metadynamics simulation before SAXS BME reweighting. (*A*) Probability distribution for the $\omega$ dihedral angle in the a03ws run 2, with the *cis*-P57 population shown. The error represents the standard deviation between the first and second halves of the analysed trajectory. (*B*) The $R_g$ probability distribution was calculated using kernel density estimates to compare the a03ws run 1 simulation (orange) and the a03ws run 2 simulation (cyan). Ensemble-averaged $R_g$ are shown for each ensemble. The associated error represents the standard deviation between the first and second halves of the analysed trajectories. The experimental SAXS data and error (standard deviation from the Guinier analysis) are shown in grey. (*C*) Ensemble-averaged, residue-specific $C^{\alpha}$ inter-residue minimum distance contact maps for a03ws run 2. The low probability of α-helix and β-sheet contacts indicates that the a03ws run 2 NAc-ORF6$_{CTR}$ conformational ensemble lacks substantial secondary structure.

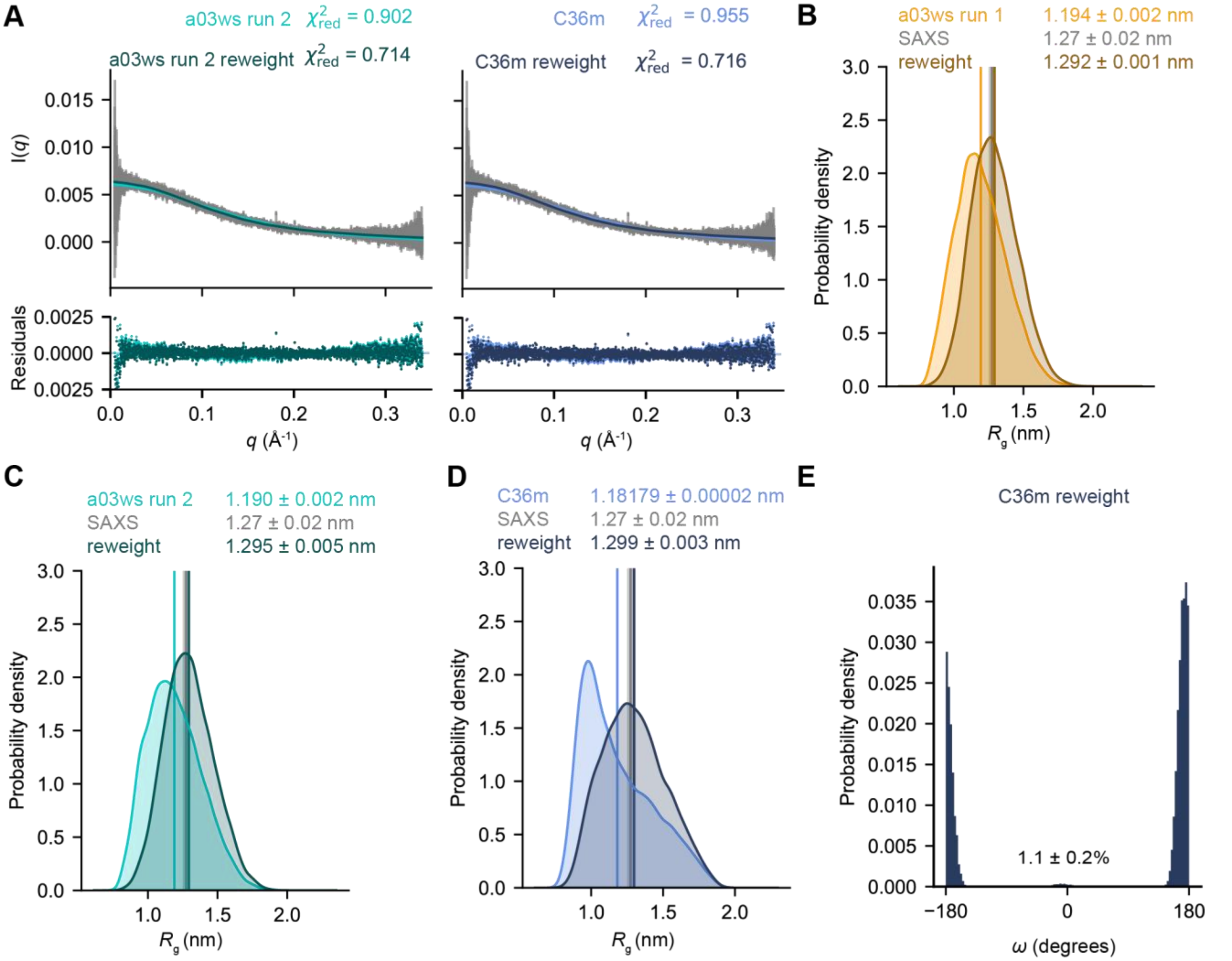


**Figure S12. NAc-ORF6$_{CTR}$ metadynamics simulations BME SAXS reweighting: a03ws run 1, a03ws run 2, and C36m.** The calculated SAXS intensities from the metadynamics simulations and the reweighted simulation are compared to the experimental SAXS intensities and the error associated with each intensity (grey) for (*A*) the a03ws run 2 (left) and the C36m (right) ensembles (43). $R_g$ probability distributions were calculated using kernel density estimates to compare the distribution before (lighter shade) and after (darker shade) the SAXS BME (44,45) reweighting for the (*B*) a03ws run 1, (*C*) a03ws run 2, and (*D*) C36m ensembles. Ensemble-averaged $R_g$ are shown for each system. The associated error represents the standard deviation between the first and second halves of the analysed trajectories. The experimental SAXS data and error (standard deviation from the Guinier analysis) are shown in grey. (*E*) Probability distribution for the $\omega$ dihedral angle in the C36m SAXS BME reweighted ensemble with the *cis*-P57 population shown. The error represents the standard deviation between the first and second halves of the analysed trajectory.

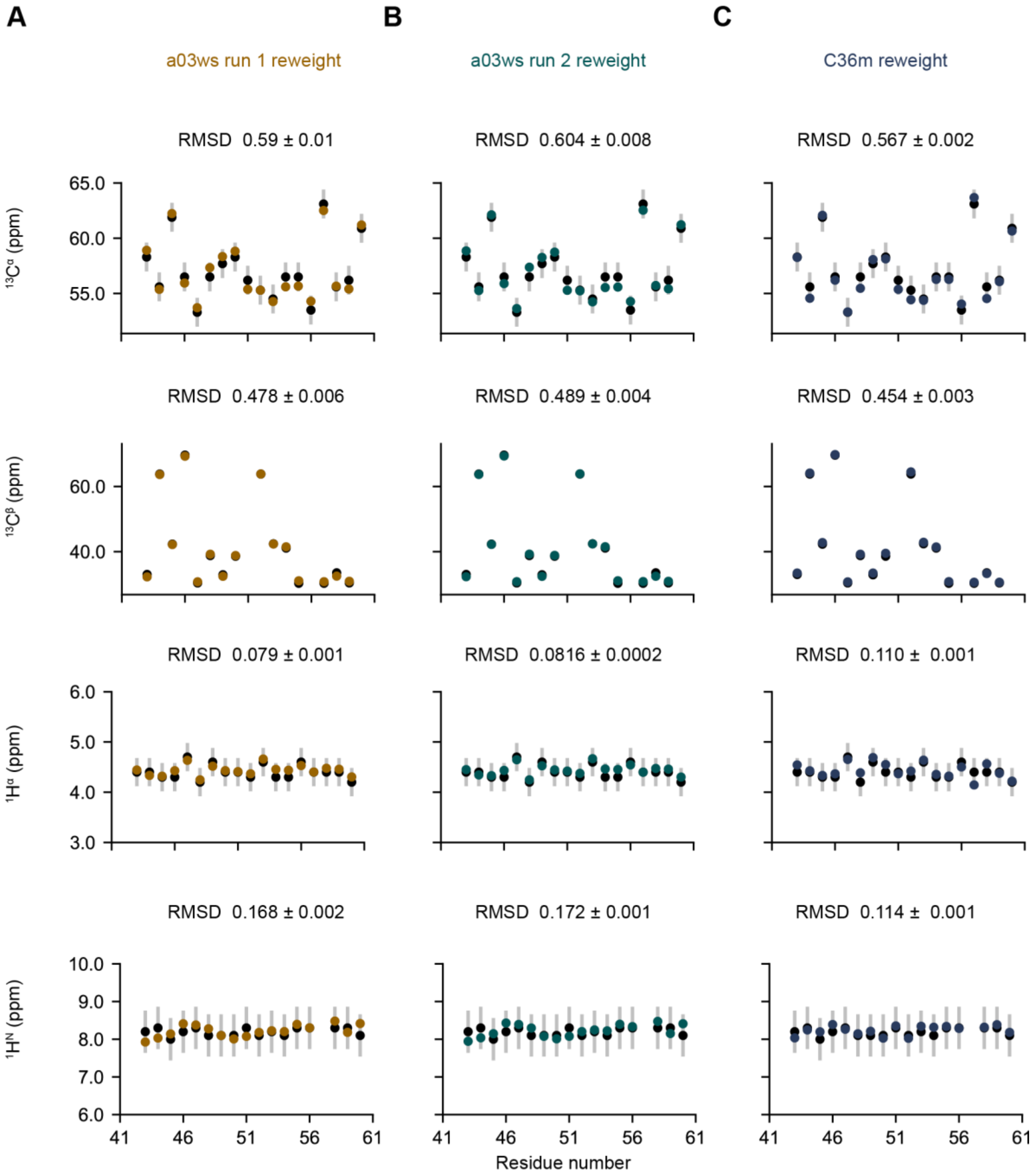


**Figure S13. Assessment of SAXS BME reweighted metadynamics simulations consistency with NMR chemical shift data: a03ws run 1, a03ws run 2, and C36m ensembles.** Comparison of the ensemble-averaged predicted chemical shifts calculated using CamShift (42) for the (*A*) a03ws run 1 (brown), (*B*) a0ws run 2 (teal), and (*C*) C36m (dark blue) ensembles to experimental chemical shifts (black) using the $^{13}C^{\alpha}$, $^{13}C^{\beta}$, $^{1}H^{\alpha}$, and $^{1}H^{N}$ chemical shifts. The error in CamShift (silver) is shown. The standard deviation between the chemical shifts predicted in the first and second halves of the analysed trajectory were also plotted but are smaller than the data points. The RMSD and its corresponding error between experimental and chemical shifts predicted from the simulations are displayed for each ensemble and chemical shift.

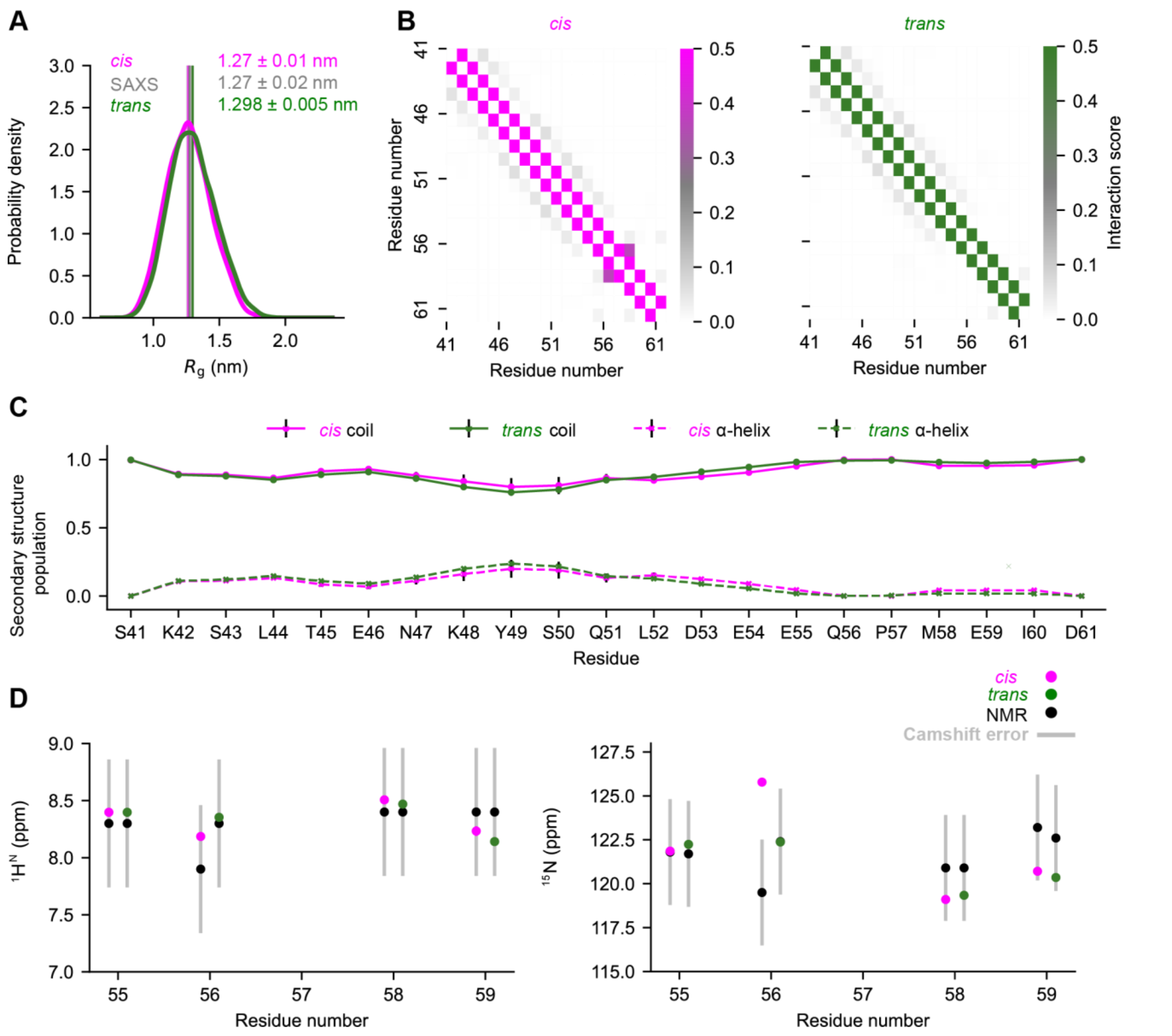


**Figure S14. NAc-ORF6$_{CTR}$ *cis*-P57 and *trans*-P57 sub-ensembles predicted by the a03ws run 2 SAXS BME reweighted metadynamics simulation have very similar profiles.** *(A)* $R_g$ probability distributions were calculated using kernel density estimates to compare the *cis*-P57 (pink) and *trans*-P57 (green) conformational sub-ensembles. Ensemble-averaged $R_g$ are shown for each conformation. The associated error represents the standard deviation between the first and second halves of the analysed trajectory. The experimental SAXS data and error (standard deviation from the Guinier analysis) are shown in grey. (*B*) C$^{\alpha}$ minimum distance contact maps for the *cis*-P57 (pink) and *trans*-P57 (green) conformations. There is a low probability of α-helix and β-strand contacts in both configurations. (*C*) Secondary structure populations for all residues in the *cis*-P57 (pink) and the *trans*-P57 (green) conformations based on SAXS BME statistical weights. Coil populations are represented by solid lines and α-helical populations by dashed lines. β-strand represents less than 0.9% of the population for each residue so was not included. Error bars (black) represent the standard deviation between the first and second halves of the analysed trajectory. (*D*) Consistency of the $^{1}H^{N}$ and $^{15}N$ *cis*-P57 and *trans*-P57 ORF6$_{CTR}$ experimental chemical shifts with the predicted chemical shifts from the SAXS BME reweighted NAc-ORF6$_{CTR}$ a03ws run 2 ensemble. The error in CamShift (silver) is shown. The standard deviation for predicted chemical shifts between the first and second halves of the analysed trajectory were also plotted but are too small to see.

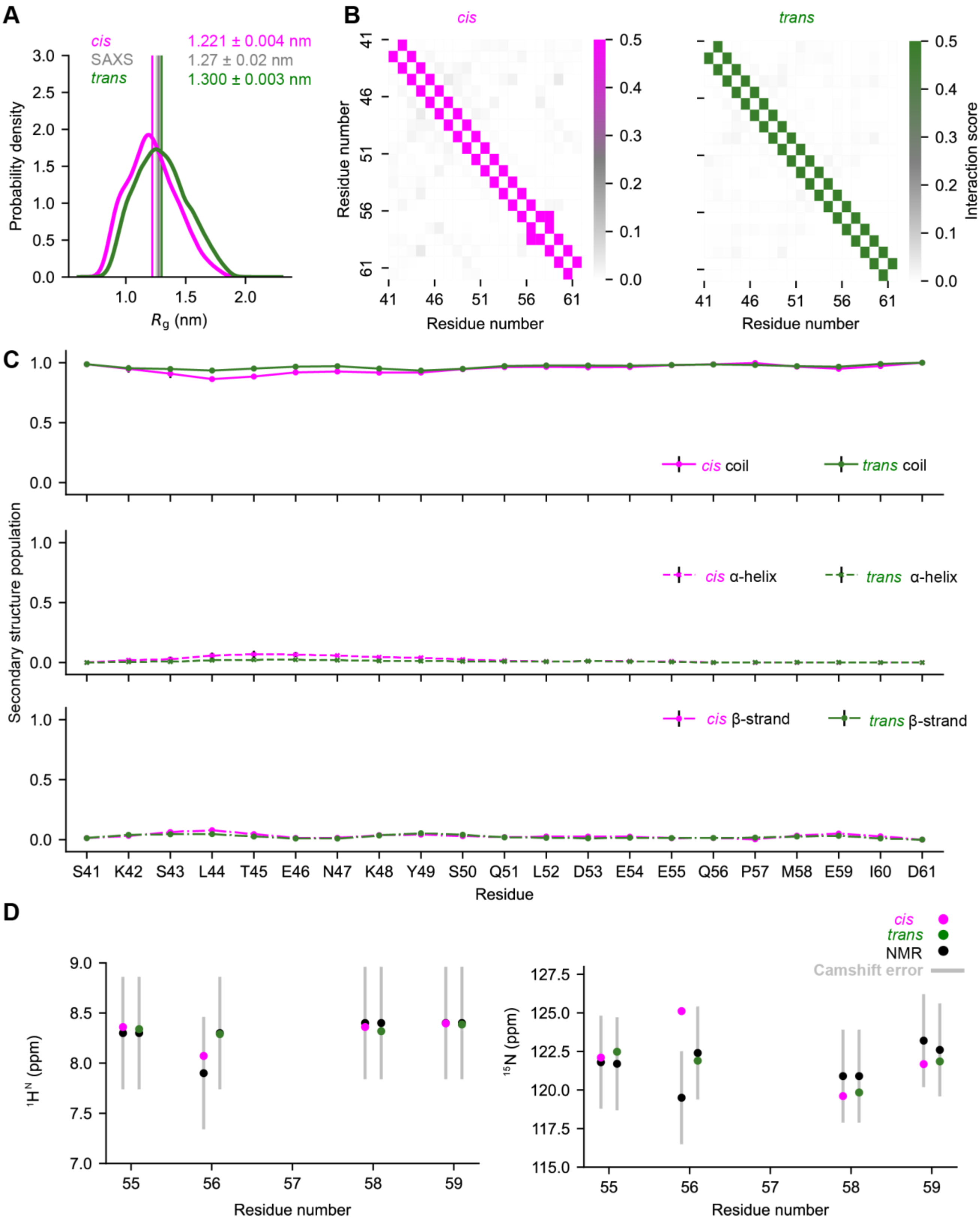
A
cis
SAXS
trans
1.221 ± 0.004 nm
1.27 ± 0.02 nm
1.300 ± 0.003 nm
Probability density
Rg (nm)
B
cis
trans
Residue number
Interaction score
C
cis coil
trans coil
cis α-helix
trans α-helix
cis β-strand
trans β-strand
Secondary structure population
S41 K42 S43 L44 T45 E46 N47 K48 Y49 S50 Q51 L52 D53 E54 E55 Q56 P57 M58 E59 I60 D61
Residue
D
cis
trans
NMR
Camshift error
1HN (ppm)
15N (ppm)
Residue number

**Figure S15. NAc-ORF6$_{CTR}$ *cis*-P57 and *trans*-P57 sub-ensembles predicted by the C36m SAXS BME reweighted metadynamics simulation have very similar profiles.** *(A)* $R_g$ probability distributions were calculated using kernel density estimates to compare the *cis*-P57 (pink) and *trans*-P57 (green) conformational sub-ensembles. Ensemble-averaged $R_g$ are shown for each conformation. The associated error represents the standard deviation between the first and second halves of the analysed trajectory. The experimental SAXS data and error (standard deviation from the Guinier analysis) are shown in grey. (*B*) $C^{\alpha}$ minimum distance contact maps for the *cis*-P57 (pink) and *trans*-P57 (green) conformations. There is a low probability of α-helix and β-strand contacts in both configurations. (*C*) Secondary structure populations for all residues in the *cis*-P57 (pink) and the *trans*-P57 (green) conformations based on SAXS BME statistical weights are shown for coil (top) α-helix (middle), and β-strand (bottom). Error bars (black) represent the standard deviation between the first and second halves of the analysed trajectory. (*D*) Consistency of the $^{1}H^{N}$ and $^{15}N$ *cis*-P57 and *trans*-P57 ORF6$_{CTR}$ experimental chemical shifts with the predicted chemical shifts from the SAXS BME reweighted NAc-ORF6$_{CTR}$ C36m ensemble. The error in CamShift (silver) is shown. The standard deviation for predicted chemical shifts between the first and second halves of the analysed trajectory were also plotted but are too small to see.

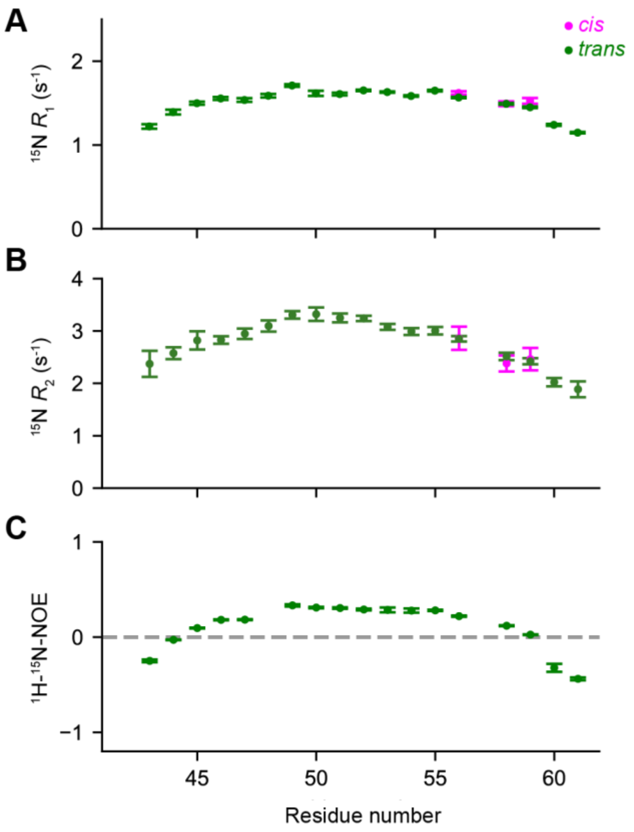


**Figure S16. The *cis*-P57 and *trans*-P57 sub-ensembles exhibit similar backbone motions at a static magnetic field strength of 18.8 T.** (*A*) $^{15}$N longitudinal relaxation rate ($R_1$), (*B*) $^{15}$N transverse relaxation rate ($R_2$) and (*C*) {$^{1}$H}-$^{15}$N nuclear Overhauser effect (hetNOE) of 300 µM $^{15}$N-labelled ORF6$_{CTR}$ at 15˚C. The backbone motions are shown for all of peaks corresponding to the *trans*-P57 conformation (green), and only for the well-resolved peaks with a measurable intensity corresponding to the *cis*-P57 (pink) conformation. Error bars represent the fitting of the NMR data in (*A*) and (*C*). The errors for (*B*) were calculated by error propagation. All experiments were recorded at a static magnetic field strength of 18.8 T and at pH 6.9.

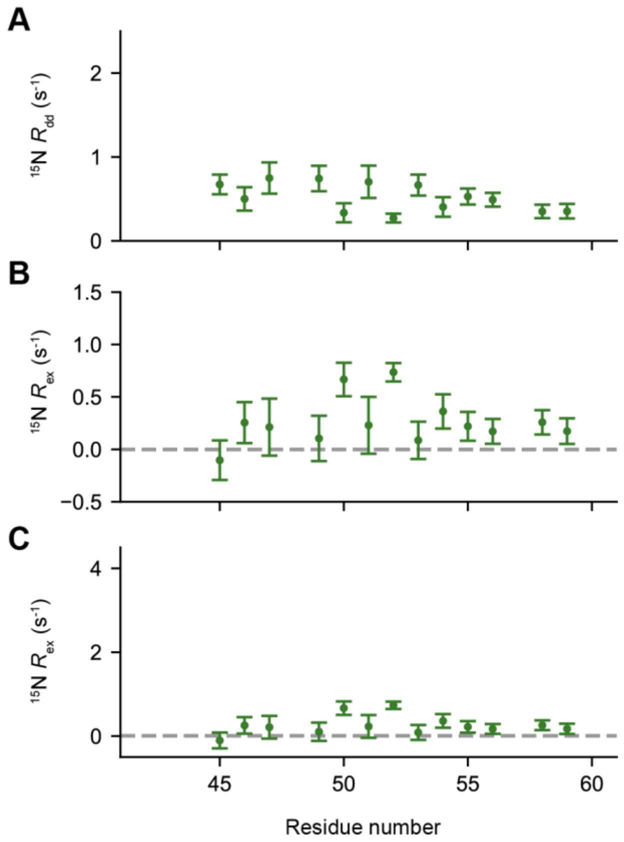


**Figure S17. Absence of microsecond to millisecond dynamics in the ORF6$_{CTR}$ *trans*-P57 conformational sub-ensemble.** (*A*) Exchange-free measure of dipole-dipole $^{15}$N transverse relaxation ($R_{dd}$) (29) , (*B*) zoomed-in view of chemical exchange analysis and (*C*) chemical exchange analysis for the *trans*-P57 conformation of 300 µM $^{15}$N-labelled ORF6$_{CTR}$ at 15˚C. The error bars in (*A*) represent the standard error associated with the exponential decay model fit for each residue. The error bars for (*B*) and (*C*) represent the standard deviations for each residue, calculated using Monte Carlo uncertainty propagation. All experiments were recorded at a static magnetic field strength of 14.1 T and at pH 6.9.

### SUPPORTING REFERENCES


1. Shaka AJ, Lee CJ, Pines A. 1988. Iterative schemes for bilinear operators; application to spin decoupling. *Journal of Magnetic Resonance*. 77:274–93.
2. Hwang TL, Shaka AJ. 1995. Water Suppression That Works. Excitation Sculpting Using Arbitrary Wave-Forms and Pulsed-Field Gradients. *J Magn Reson A*. 112:275–9.
3. Palmer AG, Cavanagh J, Wright PE, Rance M. 1991. Sensitivity improvement in proton-detected two-dimensional heteronuclear correlation NMR spectroscopy. *Journal of Magnetic Resonance*. 93:151–70.
4. Kay LE, Keifer P, Saarinen T. 1992. Pure Absorption Gradient Enhanced Heteronuclear Single Quantum Correlation Spectroscopy with Improved Sensitivity. *J Am Chem Soc*. 114:10663–5.
5. Schleucher J, Schwendinger M, Sattler M, Schmidt P, Schedletzky O, Glaser SJ, Sørensen OW, Griesinger C. 1994. A general enhancement scheme in heteronuclear multidimensional NMR employing pulsed field gradients. *J Biomol NMR*. 4:301–6.
6. Kay LE, Ikura M, Tschudin R, Bax A. 1990. Three-dimensional triple-resonance NMR spectroscopy of isotopically enriched proteins. *Journal of Magnetic Resonance*. 89:496–514.
7. Ikura M, Kay LE, Bax A. 1990. A novel approach for sequential assignment of 1H, 13C, and 15N spectra of proteins: heteronuclear triple-resonance three-dimensional NMR spectroscopy. Application to calmodulin. *Biochemistry*. 29:4659–67.
8. Clubb RT, Thanabal V, Wagner G. 1992. A constant-time three-dimensional triple-resonance pulse scheme to correlate intraresidue 1HN, 15N, and 13C′ chemical shifts in 15N-13C-labelled proteins. *Journal of Magnetic Resonance*. 97:213–7.
9. Wittekind M, Mueller L. 1993. HNCACB, a High-Sensitivity 3D NMR Experiment to Correlate Amide-Proton and Nitrogen Resonances with the Alpha- and Beta-Carbon Resonances in Proteins. *J Magn Reson B*. 101:201–5.
10. Frenkiel T, Bauer C, Carr MD, Birdsall B, Feeney J. 1990. HMQC-NOESY-HMQC, a three-dimensional NMR experiment which allows detection of nuclear overhauser effects between protons with overlapping signals. *Journal of Magnetic Resonance*. 90:420–5.
11. Ikura M, Bax A, Marius Clore G, Gronenborn AM. 1990. Detection of Nuclear Overhauser Effects between Degenerate Amide Proton Resonances by Three-Dimensional Nuclear Magnetic Resonance Spectroscopy. *J Am Chem Soc*. 112:9020–2.
12. Marion D, Driscoll PC, Kay LE, Wingfield PT, Bax A, Gronenborn AM, Clore GM. 1989. Overcoming the overlap problem in the assignment of 1H NMR spectra of larger proteins by use of three-dimensional heteronuclear 1H-15N Hartmann-Hahn-multiple quantum coherence and nuclear Overhauser-multiple quantum coherence spectroscopy: application to calmodulin. *Biochemistry*. 28:6150–6.
13. Grzesiek S, Anglister J, Bax A. 1993. Correlation of Backbone Amide and Aliphatic Side-Chain Resonances in 13C/15N-Enriched Proteins by Isotropic Mixing of 13C Magnetization. *J Magn Reson B*. 101:114–9.
14. Kay LE, Xu GY, Singer AU, Muhandiram DR, Formankay JD. 1993. A Gradient-Enhanced HCCH-TOCSY Experiment for Recording Side-Chain 1H and 13C Correlations in H2O Samples of Proteins. *J Magn Reson B*. 101:333–7.
15. Bax A, Clore GM, Gronenborn AM. 1990. 1H-1H correlation via isotropic mixing of 13C magnetization, a new three-dimensional approach for assigning 1H and 13C spectra of 13C-enriched proteins. *Journal of Magnetic Resonance*. 88:425–31.
16. Bermel W, Bertini I, Duma L, Felli IC, Emsley L, Pierattelli R, Vasos PR. 2005. Complete Assignment of Heteronuclear Protein Resonances by Protonless NMR Spectroscopy. *Angewandte Chemie International Edition*. 44:3089–92.
17. Duma L, Hediger S, Lesage A, Emsley L. 2003. Spin-state selection in solid-state NMR. *Journal of Magnetic Resonance*. 164:187–95.
18. Kay LE, Xu GY, Yamazaki T. 1994. Enhanced-Sensitivity Triple-Resonance Spectroscopy with Minimal H2O Saturation. *J Magn Reson A*. 109:129–33.

19. Muhandiram DR, Kay LE. 1994. Gradient-Enhanced Triple-Resonance Three-Dimensional NMR Experiments with Improved Sensitivity. *J Magn Reson B*. 103:203–16.
20. Kadkhodaie M, Rivas O, Tan M, Mohebbi A, Shaka AJ. 1991. Broadband homonuclear cross polarization using flip-flop spectroscopy. *Journal of Magnetic Resonance*. 91:437–43.
21. Kay LE, Torchia DA, Bax A. 1989. Backbone Dynamics of Proteins As Studied by 15N Inverse Detected Heteronuclear NMR Spectroscopy: Application to Staphylococcal Nuclease. *Biochemistry*. 28:8972–9.
22. Dayie KT, Wagner G. 1994. Relaxation-Rate Measurements for 15N−1H Groups with Pulsed-Field Gradients and Preservation of Coherence Pathways. *J Magn Reson A*. 111:121–6.
23. Kay LE, Nicholson LK, Delaglio F, Bax A, Torchia DA. 1992. Pulse sequences for removal of the effects of cross correlation between dipolar and chemical-shift anisotropy relaxation mechanisms on the measurement of heteronuclear T1 and T2 values in proteins. *Journal of Magnetic Resonance*. 97:359–75.
24. Farrow NA, Muhandiram R, Pascal SM, Kay LE, Singer AU, Forman-Kay JD, et al. 1994. Backbone dynamics of a free and phosphopeptide-complexed Src homology 2 domain studied by 15N NMR relaxation. *Biochemistry*. 33:5984–6003.
25. Korzhnev DM, Skrynnikov NR, Millet O, Torchia DA, Kay LE. 2002. An NMR experiment for the accurate measurement of heteronuclear spin-lock relaxation rates. *J Am Chem Soc*. 124:10743–53.
26. Hansen DF, Kay LE. 2007. Improved magnetization alignment schemes for spin-lock relaxation experiments. *J Biomol NMR*. 37:245–55.
27. Ferrage F, Cowburn D, Ghose R. 2009. Accurate sampling of high-frequency motions in proteins by steady-state 15N-{ 1H} nuclear overhauser effect measurements in the presence of cross-correlated relaxation. *J Am Chem Soc*. 131:6048–9.
28. Li YC, Montelione GT. 1994. Overcoming solvent saturation-transfer artifacts in protein NMR at neutral pH. Application of pulsed field gradients in measurements of 1H-15N Overhauser effects. *J Magn Reson B*. 105:45–51.
29. Hansen DF, Yang D, Feng H, Zhou Z, Wiesner S, Bai Y, Kay LE. 2007. An exchange-free measure of 15N transverse relaxation: an NMR spectroscopy application to the study of a folding intermediate with pervasive chemical exchange. *J Am Chem Soc*. 129:11468–79.
30. Wu DH, Chen A, Johnson CS. 1995. An Improved Diffusion-Ordered Spectroscopy Experiment Incorporating Bipolar-Gradient Pulses. *J Magn Reson A*. 115:260–4.
31. Stejskal EO, Tanner JE. 1965. Spin Diffusion Measurements: Spin Echoes in the Presence of a Time-Dependent Field Gradient. *J Chem Phys*. 42:288–92.
32. Bonomi M, Bussi G, Camilloni C, Tribello GA, Banáš P, Barducci A, et al. 2019. Promoting transparency and reproducibility in enhanced molecular simulations. *Nat Methods*. 16:670–3.
33. Heller GT, Aprile FA, Michaels TCT, Limbocker R, Perni M, Ruggeri FS, et al. 2020. Small-molecule sequestration of amyloid-β as a drug discovery strategy for Alzheimer’s disease. *Sci Adv*. 6.
34. Pietrucci F, Laio A. 2009. A collective variable for the efficient exploration of protein beta-sheet structures: Application to SH3 and GB1. *J Chem Theory Comput*. 5:2197–201.
35. Melis C, Bussi G, Lummis SCR, Molteni C. 2009. Trans-cis switching mechanisms in proline analogues and their relevance for the gating of the 5-HT3 receptor. *J Phys Chem B*. 113:12148–53.
36. Maschio MC, Fregoni J, Molteni C, Corni S. 2021. Proline isomerization effects in the amyloidogenic protein β 2 -microglobulin. *Physical Chemistry Chemical Physics*. 23:356–67.
37. Pfaendtner J, Bonomi M. 2015. Efficient Sampling of High-Dimensional Free-Energy Landscapes with Parallel Bias Metadynamics. *J Chem Theory Comput*. 11:5062–7.
38. Sormanni P, Camilloni C, Fariselli P, Vendruscolo M. 2015. The s2D Method: Simultaneous Sequence-Based Prediction of the Statistical Populations of Ordered and Disordered Regions in Proteins. *J Mol Biol*. 427:982–96.
39. Jumper J, Evans R, Pritzel A, Green T, Figurnov M, Ronneberger O, et al. 2021. Highly accurate protein structure prediction with AlphaFold. *Nature*. 596:583–9.
40. Pesce F, Newcombe EA, Seiffert P, Tranchant EE, Olsen JG, Grace CR, Kragelund BB, Lindorff-Larsen K. 2023. Assessment of models for calculating the hydrodynamic radius of intrinsically disordered proteins. *Biophys J*. 122:310–21.

41. Bussi G, Tribello GA. 2019. Analyzing and Biasing Simulations with PLUMED. *Methods in molecular biology*. 2022:529–78.
42. Kohlhoff KJ, Robustelli P, Cavalli A, Salvatella X, Vendruscolo M. 2009. Fast and accurate predictions of protein NMR chemical shifts from interatomic distances. *J Am Chem Soc*. 131:13894–5.
43. Grudinin S, Garkavenko M, Kazennov A. 2017. Pepsi-SAXS: an adaptive method for rapid and accurate computation of small-angle X-ray scattering profiles. *Acta Crystallogr D Struct Biol*. 73:449–64.
44. Bottaro S, Bengtsen T, Lindorff-Larsen K. 2020. Integrating Molecular Simulation and Experimental Data: A Bayesian/Maximum Entropy Reweighting Approach. *Methods in molecular biology*. 2112:219–40.
45. Ahmed MC, Skaanning LK, Jussupow A, Newcombe EA, Kragelund BB, Camilloni C, Langkilde AE, Lindorff-Larsen K. 2021. Refinement of α-Synuclein Ensembles Against SAXS Data: Comparison of Force Fields and Methods. *Front Mol Biosci*. 8:654333.
